# Species-specific basal fluidization shapes early forebrain development

**DOI:** 10.1101/2025.10.07.680907

**Authors:** Shuting Xu, Sangwoo Kim, Guanlin Li, Michaela Wilsch-Bräuninger, Annika Kolodziejczyk, Theresa M. Schütze, Mareike Albert, Vikas Trivedi, Otger Campàs, Miki Ebisuya

## Abstract

The human forebrain exhibits an expanded surface area compared with other great apes. Despite progress in dissecting species-specific gene expression programs and cell lineages, the tissue-level physical mechanisms underlying human-specific neurodevelopment remain unclear. Here, by quantifying tissue dynamics and mechanics using oil microdroplets in cerebral organoids from humans, gorillas, chimpanzees, and mice, we uncover a transient tissue fluidization occurring exclusively in the basal region of human neuroepithelia. Enhanced droplet motility and frequent cell rearrangements reveal this basal fluidization, which gradually diminishes at the onset of neurogenesis in humans. By contrast, such basal fluidization is not detected in non-human neuroepithelia examined here. Mechanistically, basal fluidization is driven by basal nuclear fluctuations and facilitated by expanded intercellular spaces. Increasing N-cadherin expression in human neuroepithelia to gorilla levels suppresses both nuclear fluctuations and basal fluidization. Integrating simulations of neuroepithelial dynamics with experimental data, we further find that nuclear fluctuations underlying basal fluidization promote the insertion of newly divided nuclei into the basal side to support tangential surface expansion, a hallmark of human forebrain development.

## Introduction

Among primates, the human brain is distinguished by its exceptional size in both thickness and surface area. The human neocortex alone is three times larger than that of non-human great apes and contains roughly twice as many cortical neurons^1–4^. This evolutionary expansion, particularly the tangential (lateral) expansion of the human forebrain surface, begins prior to neurogenesis and is initially driven by more active symmetric divisions of early neural progenitors, known as neuroepithelial cells^5–7^. However, comprehensive mechanisms underlying species-specific forebrain expansion remain unclear, as direct examination of early forebrain development in human and great ape embryos is ethically precluded, making alternative approaches necessary.

Recent advances in stem-cell-derived brain organoids have opened a research window into early human forebrain development^8–11^. The generation of brain organoids from great apes has further enabled cross-species comparisons in vitro. Comparative brain organoid studies, combined with single-cell genomics and genetic manipulation, have uncovered human-specific gene expression patterns, distinct cell types, and a unique timeline of human neurodevelopment^4,5,8,12–19^. Cross-species genomic analyses have also highlighted the contributions of human-specific genes, cis-regulatory elements, and genomic variants^2,3,20,21^. However, to fully understand how interspecies differences at the molecular and cellular levels give rise to organ-level features, it is essential to investigate the tissue dynamics and mechanics that shape brain development across species.

In contrast to molecular and cellular insights, the role of tissue mechanics in brain development remains less explored. Substrate stiffness has been shown to influence neural progenitor lineage choice, and cortical folding during late fetal stages is driven by mechanical compression^22–24^. However, systematic comparisons between humans and non-human great apes are lacking, largely due to the difficulty of probing mechanics in developing brain tissues. Techniques such as atomic force microscopy and micropipette aspiration are primarily restricted to tissue surfaces, while Brillouin microscopy requires prior knowledge of tissue optical properties and density^25–28^. Injecting fluorescent microdroplets and quantifying their deformation and dynamics provides an effective means of examining mechanics inside complex 3D tissues^29,30^. In this study, we applied the microdroplet approach to cerebral organoids derived from humans and other species, and identified basal tissue fluidization as a mechanical hallmark of human neuroepithelia. Combining simulations of pseudostratified neuroepithelia with experimental data, we show that basal nuclear fluctuations promote basal fluidization and the nuclear insertion required for tangential surface expansion.

## Results

### Neuroepithelia exhibit transiently enhanced tissue dynamics in humans, but not in non-human apes or mice

We generated cerebral organoids from human induced pluripotent stem cells (iPSCs) (IMR90-4 line), gorilla iPSCs, and mouse embryonic stem cells (ESCs) (Figure 1, A and B; Figure S1A; Methods). Gorillas and mice were selected to represent non-human great apes and non-primate mammals, respectively, as their cerebral organoids have been characterized using protocols comparable to those for human organoids^5,11,31^. Gorilla cerebral organoids, in particular, have been reported to exhibit developmental timings more similar to humans than to chimpanzees^5^. Our day numbering differs from the previously reported protocol due to a different definition of the starting day^5^. Human and gorilla organoids indeed formed distinct neural buds of pseudostratified neuroepithelium by day 10 following Matrigel-induced epithelialization, whereas mouse organoids developed more rapidly, forming buds by day 6.5 (Figure 1C; Figure S1B). Neurogenesis began between days 15-20 in human and gorilla organoids, whereas it was initiated earlier, around days 6.5-7, in mouse organoids^5^. Among the three species, human organoids exhibited the largest neural bud perimeter, indicating that human-specific tangential expansion initiates at the neuroepithelial stage, consistent with the previous report^5^ (Figure 1D). Despite variability across organoids, the interspecies differences remained robust.

**Figure 1.**
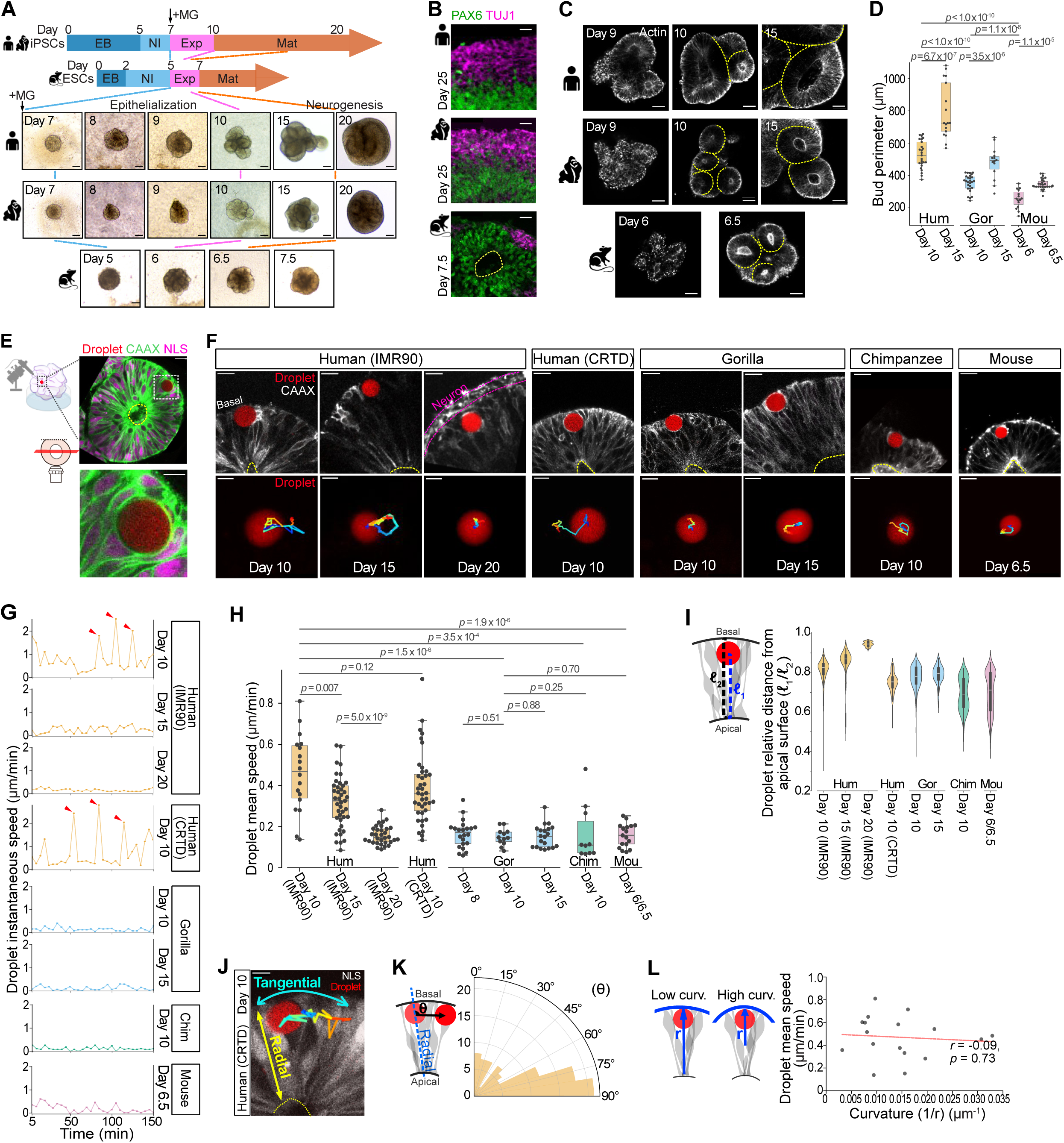
Human neuroepithelia exhibit unique tissue dynamics. (A) Protocols and bright-field images of cerebral organoids derived from human, gorilla, and mouse cells. Dashed lines indicate corresponding stages across species. iPSCs, induced pluripotent stem cells; ESCs, embryonic stem cells; EB, embryoid body; NI, neural induction; Exp, neuroepithelial expansion; Mat, maturation; MG: Matrigel. Scale bar: 100 μm. (B) Immunofluorescence images showing neural progenitor marker PAX6 (green) and neuronal marker TUJ1 (magenta) in human and gorilla organoids at day 25, and mouse organoid at day 7.5. Yellow dashed line marks the apical surface. Scale bar: 20 μm. (C) SiR-actin labeling of neural bud morphology transformations during organoid epithelization. Yellow dashed lines outline individual neural buds. Hum, human; Gor, gorilla; Mou, mouse. Scale bar: 50 μm. (D) Neural bud outer perimeters (2D surface outlines) across species and developmental stages. Human day 10 (n = 25 neural buds, N = 6 independent organoid batches), human day 15 (n = 17, N = 6), gorilla day 10 (n = 27, N = 3), gorilla day 15 (n = 14, N = 3), mouse day 6 (n = 15, N = 2), mouse day 6.5 (n = 29, N = 2). (E) Schematic (left) and representative image (right) of the oil droplet assay in cerebral organoid expressing EGFP-CAAX (membrane, green) and mCherry-NLS (nuclei, magenta). Scale bar: 20 μm (top), 10 μm (bottom). (F) Droplets positioned in the basal region of organoids across species and stages (top) with corresponding color-coded trajectories over a 3-h period (bottom). Membranes are labeled by EGFP-CAAX (white), and the droplet is shown in red. Trajectories start in blue and end in red. Yellow dashed lines mark the apical surface. Scale bar: 20 μm (top), 10 μm (bottom). (G) Representative traces of droplet instantaneous speed during 150-min imaging across species and stages. Red arrowheads highlight large-displacement events (“jumps”) observed in day 10 human organoids. (H) Droplet mean speed. Each data point represents the mean speed of one droplet. Mouse days 6 and 6.5 data were pooled, as in Figure S1, C and G. Human IMR90 day 10 (n = 18 droplets), human IMR90 day 15 (n = 41), human IMR90 day 20 (n = 32), human CRTD day 10 (n = 40), gorilla day 8 (n = 22), gorilla day 10 (n = 12), gorilla day 15 (n = 20), chimpanzee day 10 (n = 10), mouse days 6/6.5 (n = 17). (I) Droplet positions relative to the apical surface. ℓ[ denotes the distance from the droplet centroid to the apical surface; ℓ[ indicates the total tissue thickness. In day 20 human organoids, ℓ[ corresponds to the thickness of the ventricular zone, excluding the neural layer. Each data point corresponds to one droplet at one time point. Human IMR90 day 10 (n = 561 data points, N =15 droplets), human day 15 (n = 517, N = 14), human day 20 (n = 398, N = 13), human CRTD day 10 (n = 480, N =20), gorilla day 10 (n = 260, N = 12), gorilla day 15 (n = 420, N = 15), chimpanzee day 10 (n = 208, N = 8), mouse days 6/6.5 (n = 362, N = 14). (J) Droplet moves tangentially, perpendicular to the apical-basal radial axis, in a day 10 human organoid. The trace spans 155 min. Scale bar: 20 μm. (K) Polar histogram of droplet movement angles (θ). θ is defined as the angle between the line connecting droplet centroids over four time points (t, t+4) and the local radial axis at t; values near 90° indicate tangential orientation. n = 93 calculations from 16 droplets. Bin size is 10. (L) Relationship between local curvature (i.e., inverse radius) of basal neuroepithelial surface and droplet mean speed in day 10 human organoids. Red line shows linear regression fit; r and p-values from Pearson correlation. n = 17 droplets.

To probe tissue mechanics and dynamics potentially contributing to species-specific forebrain development, we injected fluorescent oil microdroplets into the intercellular spaces of neural buds^29,32,33^ (Figure 1E). The droplets, 7-15 μm in radius and larger than individual nuclei (Figure 1E; Figure S1C), were imaged after at least 5 h of post-injection recovery. Unexpectedly, droplets moved around within human neuroepithelia on day 10, occasionally displaying long-range displacements, as indicated by their speed (Figure 1, F and G; Figure S1D; Video S1). This droplet motility declined by day 15 and largely ceased by day 20, coinciding with the onset of neurogenesis (Figure 1H). By contrast, droplet motility was negligible in gorilla neuroepithelia on day 10 and at later stages, and similarly minimal in mouse neuroepithelia on days 6-6.5 (Figure 1, F to H; Figure S1D; Video S2).

The absence of droplet motility in gorilla and mouse organoids cannot be attributed to missing the optimal time window, as day 10 in gorillas and day 6 in mice are the earliest stages at which individual neural buds become morphologically distinguishable (Figure S1B; Figure 1C). Even when droplets were injected into gorilla organoids on day 8, corresponding to premature neural buds with barely formed lumens or to aggregates lacking discernible bud structures (Figure S1E), no droplet motility was observed (Figure 1H; Figure S1D). Thus, despite the faster neurogenesis in mouse organoids and the potentially faster development of gorilla organoids relative to humans (although neurogenesis timing in gorillas has been reported to resemble that of humans^5^), transient droplet motility appears to be specific to human organoids. To exclude cell line-specific effects, we used an independent human iPSC line (CRTDi011-A) (Figure S1F) and confirmed high droplet motility in human neuroepithelia on day 10, consistent with observations from the original human iPSC line (IMR90-4) (Figure 1, F to H; Figure S1D; Video S2). Due to the limited availability of independent gorilla iPSC lines, we instead used a chimpanzee iPSC line (Figure S1F) and confirmed low droplet motility in chimpanzee neuroepithelia on day 10, comparable to that observed in gorillas (Figure 1, F to H; Figure S1D; Video S2).

Notably, droplets consistently localized to the basal region of neural buds, regardless of the initial injection sites (Figure 1I; Figure S1G), and their movement was confined to tangential trajectories oriented perpendicular to the apical-basal (radial) axis (Figure 1, J and K; Video S1). Although neural buds in organoids exhibit higher curvature than the embryonic neural tube in vivo, curvature did not significantly influence droplet motility (Figure 1L; Figure S1H). Together, these findings identify species-and stage-specific droplet motility during forebrain development, suggesting unique tissue dynamics and mechanics in the basal region of human neuroepithelia. Because gorillas and humans are phylogenetically close and exhibit comparable organoid development^5^, our subsequent analyses focused on these two species.

### Fluctuations in radial nuclear movements in the basal region correlate with tissue dynamics

To identify the driving force underlying tangential droplet movement, we examined nuclear dynamics, as nuclei are known to undergo active interkinetic nuclear migration (IKNM) along the radial axis during the cell cycle^7,34–36^ (Figure 2A). As droplets preferentially localized to the basal region (Figure 1I), we focused our analysis on nuclear movement in this region. In human neuroepithelia on day 10, nuclei in the basal region displayed fluctuations in radial movement and occasionally underwent long-range displacements in random directions (Figure 2B; Figure S2, A and B). These basal nuclear fluctuations declined by day 15 and largely ceased by day 20 (Figure 2, B to D). Furthermore, basal nuclear fluctuations were absent in gorilla organoids even on day 10 (Figure 2, B to D; Figure S2, A and B; Video S3). These patterns of nuclear fluctuations closely mirrored those of droplet motility (compare Figure 1H and Figure 2C), suggesting that nuclear dynamics in the basal region may underlie tangential droplet movement. The observed non-directional fluctuations in nuclear movements were specific to the basal region: in the apical region, nuclei directionally migrated with minimal fluctuations (Figure 2E; Figure S2, C and D), indicating a basal-to-apical gradient in nuclear fluctuation dynamics (Figure 2F).

**Figure 2.**
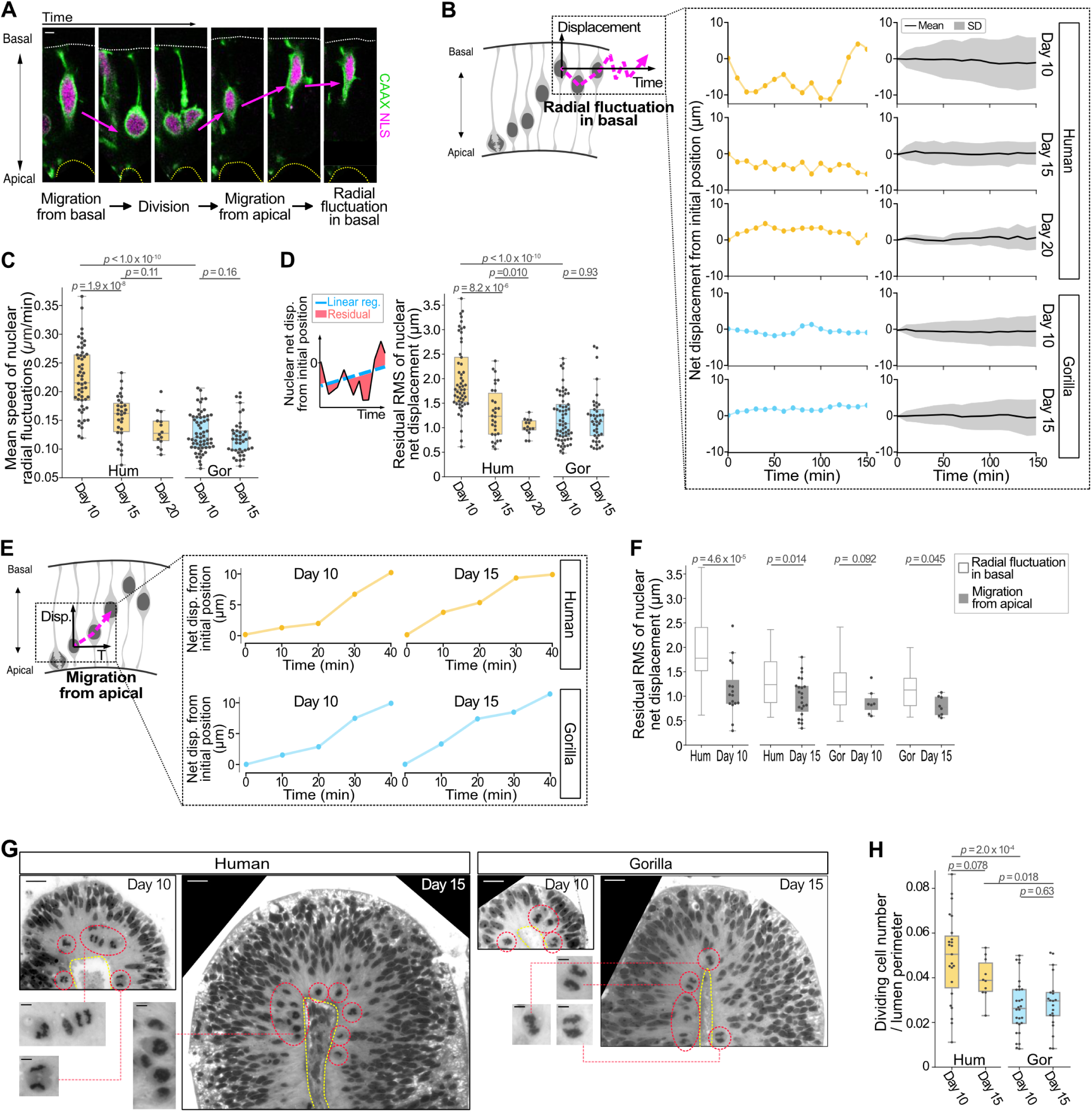
Radial nuclear fluctuations in the basal region correlate with tissue dynamics (A) Time-lapse images of interkinetic nuclear migration (IKNM). Nuclei and membranes are sparsely labeled with mCherry-NLS (magenta) and EGFP-CAAX (green), showing migration from the basal surface, division at the apical surface, migration from the apical surface, and fluctuations in radial movements in the basal region. Scale bar: 5 μm. (B) Schematic of radial fluctuations of basal nuclei (left), representative 150-min traces of nuclear net displacement from the initial position at t = 0 (middle), and mean ± SD (right) in human and gorilla organoids at the indicated days. Negative values indicate movement toward the apical surface relative to the initial position, whereas positive values indicate movement toward the basal surface. Human day 10 (n = 53 nuclei), human day 15 (n = 30), human day 20 (n = 14), gorilla day 10 (n = 61), gorilla day 15 (n = 41). (C) Mean speed of radial nuclear fluctuations across species and stages. Data as in (B, right). (D) Root-mean-square error (RMSE) of nuclear net displacement during radial fluctuations. RMSE was obtained by fitting nuclear displacement traces with a linear regression and calculating residuals between observed and predicted values. Higher RMSE indicates greater variability in nuclear displacement dynamics, reflecting stronger nuclear fluctuations. Data as in (B). (E) Schematic of nuclear migration from the apical surface (left) and representative 40-min displacement traces (right) across species and stages. Displacement is plotted as described in (B). Analysis was restricted to nuclei within 30 μm of the apical surface, rather than full apical-basal migration. (F) RMSE of nuclear net displacement during migration from the apical surface. Residuals were calculated as the difference between observed and linear-regression-predicted values. Migration from the apical surface was analyzed for nuclei located within 30 μm of the apical surface, thus reflecting apical-region tissue dynamics (migration from apical). RMSE of radial fluctuations in the basal region from (D) is also shown for comparison (fluctuation in basal). Human day 10 (n = 16 nuclei), human day 15 (n = 23), gorilla day 10 (n = 7); gorilla day 15 (n = 9). (G) Mitotic figures identified in light microscopy images of human and gorilla organoids at days 10 and 15. Red circles indicate dividing cells. Yellow outlines mark the apical surfaces. Scale bar: 20 μm (main), 5 μm (inset). (H) Number of dividing cells normalized to the lumen perimeter. Human day 10 (n = 23 neural buds), human day 15 (n = 10), gorilla day 10 (n = 28), gorilla day 15 (n = 21).

In addition to nuclear fluctuations within the basal region, droplet motility may also be influenced by the frequency of nuclei entering the basal region after division in the apical region and colliding with droplets. To test this, we quantified dividing cells near the apical surface (Figure 2G; Figure S2E). Cell division rates in human organoids were consistently higher than gorilla organoids across stages (Figure 2H), largely mirroring the patterns of droplet motility (Figure 1H). Together, these results highlight nucleus-droplet interactions as a potential driving force of droplet motility.

### Reducing nuclear fluctuations suppresses tissue dynamics

To directly assess the role of basal nuclear fluctuations in driving droplet movement, we perturbed nuclear dynamics. Treatment with the DNA replication inhibitors hydroxyurea (Hu) and aphidicolin (Aph), which block both cell cycle progression and IKNM in neural progenitors^36,37^ (Figure 3A), substantially reduced radial nuclear fluctuations in the basal region (Figure 3, B to D; Figure S3, A and B; Video S4). Notably, tangential droplet movement was also markedly suppressed, reinforcing the mechanistic link between nuclear dynamics and droplet motility (Figure 3, E and F; Figure S3, C and D; Video S4).

**Figure 3.**
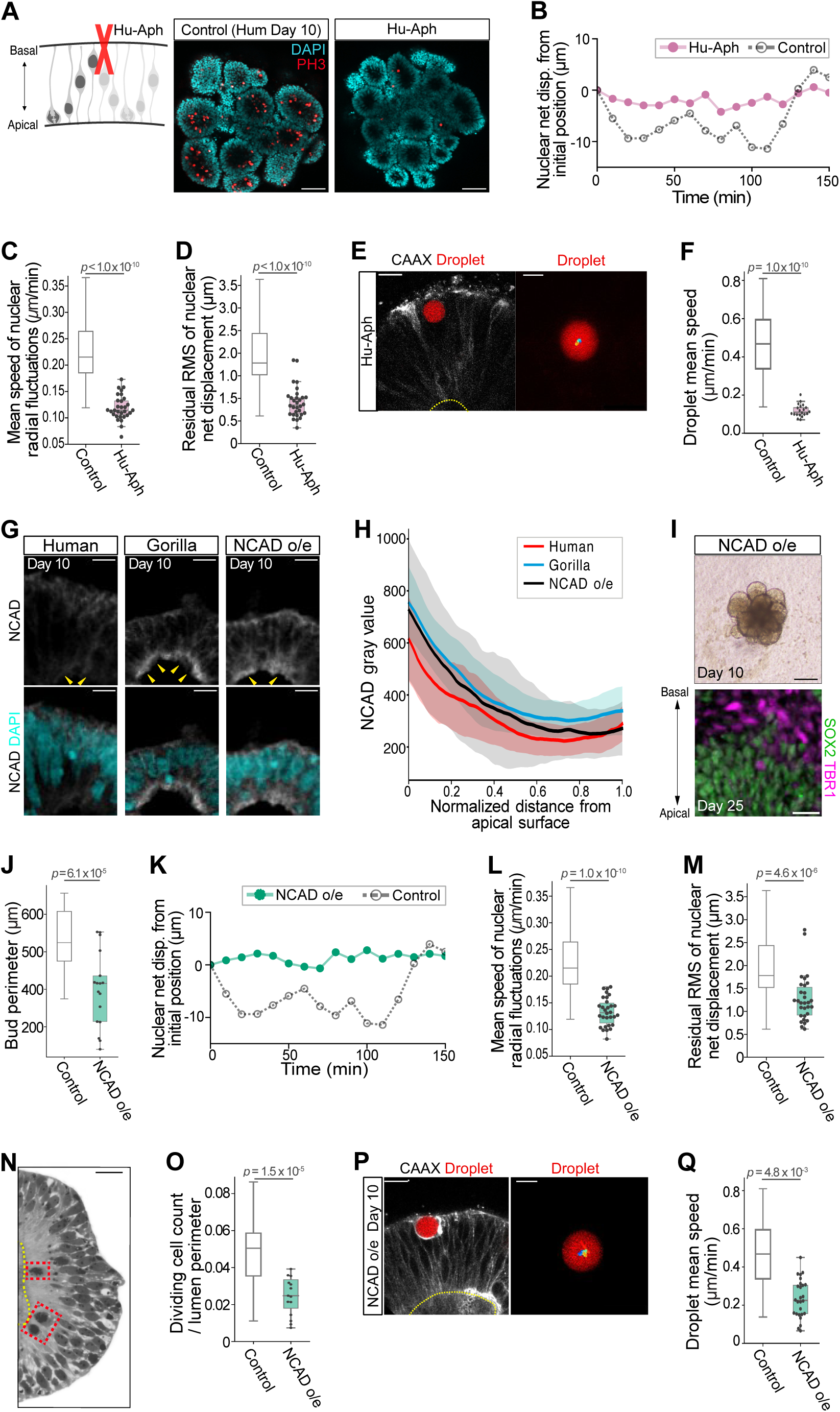
Radial nuclear fluctuations in the basal region drive tissue dynamics (A) Schematic of human organoids treated with S-phase inhibitors hydroxyurea and aphidicolin (Hu-Aph, left) and representative images (right) showing reduced mitotic cells labeled with phospho-histone 3 (PH3, red). Nuclei are labeled with DAPI (cyan). Scale bar: 50 μm. (B) Representative displacement trace of radial nuclear fluctuations in Hu-Aph-treated day 10 human organoids. Displacement is plotted as in Figure 2B. Control is from Figure 2B (human day 10). (C) Mean speed of radial nuclear fluctuations in Hu-Aph-treated organoids. Control is from Figure 2C (human day 10). n = 30 nuclei. (D) RMSE of nuclear net displacement during radial fluctuations in Hu-Aph-treated organoids. Control is from Figure 2D (human day 10). Data as in (C). (E) Droplet in a Hu-Aph-treated day 10 human organoid (left) and the corresponding color-coded trajectory over a 3-h trace (right). Droplet in red; membranes labeled with EGFP-CAAX (white); apical surface marked by yellow dashed line. Scale bar: 20 μm (left), 10 μm (right). (F) Droplet mean speed in Hu-Aph-treated organoids. Control is from Figure 1H (human day 10). n = 20 droplets. (G) N-Cadherin (NCAD, white) staining in human, gorilla, and NCAD-overexpressing human (NCAD o/e) organoids at day 10. Nuclei are labeled with DAPI (cyan). Yellow arrowheads indicate apical enrichment of NCAD. Scale bar: 10 μm. (H) NCAD intensity profiles along the apical-basal axis in human, gorilla, and NCAD o/e human organoids at day 10. X = 0 and 1 mark the apical and basal surfaces, respectively. Lines and shaded regions represent mean ± SD. Human (n = 42 neural buds), gorilla (n = 28), NCAD o/e (n = 28). (I) Bright-field (top, day 10) and immunofluorescence (bottom, day 25) images of NCAD o/e human organoids. SOX2 (green) marks neural progenitors and TBR1 (magenta) marks neurons. Scale bar: 100 μm (top), 20 μm (bottom). (J) Neural bud perimeter in day 10 NCAD o/e organoids. Control is from Figure 1D (human day 10). n = 17 neural buds. (K) Representative displacement traces of nuclear radial fluctuations in day 10 NCAD o/e organoids. Control is from Figure 2B (human day 10). (L) Mean speed of nuclear radial fluctuations in NCAD o/e organoids. Control is from Figure 2C (human day 10). n = 33 nuclei. (M) RMSE of nuclear net displacement during radial fluctuations in NCAD o/e organoids. Control is from Figure 2D (human day 10). Data as in (L). (N) Mitotic figures (red boxes) identified in light microscopy images of a day 10 NCAD o/e organoid. Yellow outlines mark the apical surface. Scale bar: 20 μm. (O) Number of dividing cells normalized to the lumen perimeter in NCAD o/e organoids. Control is from Figure 2H (human day 10). n = 14 neural buds. (P) Droplet in a day 10 NCAD o/e organoid (left) and the corresponding color-coded trajectory over a 3-h trace (right). Details and scale bar as in (E). (Q) Droplet mean speed in NCAD o/e organoids. Control is from Figure 1H (human day 10). n = 26 droplets.

In parallel with pharmacological perturbation of cell cycle and nuclear dynamics, which could potentially cause undesirable side effects, we pursued a complementary genetic approach. We focused on adhesion and tight junction molecules to restrict cell movement^38^. Among these, N-cadherin (NCAD) is known to increase as neural progenitors mature^5,39,40^, and its expression was higher in gorilla neuroepithelia on day 10 compared with human (Figure 3, G and H; Figure S3E). We therefore constitutively overexpressed NCAD in human organoids, achieving expression levels comparable to those in gorillas (Figure 3, G and H; Figure S3E). NCAD-overexpressing human organoids appeared developmentally normal but formed smaller neural buds as seen in gorillas (Figure 3, I and J). Importantly, NCAD-overexpressing organoids exhibited reduced radial nuclear fluctuations (Figure 3, K to M; Figure S3, F and G; Video S4), lower cell division rates (Figure 3, N and O), and marked suppression of tangential droplet movement (Figure 3, P and Q; Figure S3, H and I; Video S4). Together, these perturbation experiments suggest that species-and stage-specific nuclear fluctuations underlie droplet motility and thereby tissue dynamics.

### Abundant basal intercellular spaces characterize a permissive environment in human neuroepithelia

In addition to nuclear dynamics, tissue architecture and cell connectivity are likely to influence tissue dynamics and mechanics^30,38,41–43^. As droplets reside within spaces between cells in the basal region, we characterized these intercellular spaces. Transmission electron microscopy (TEM) imaging revealed that the basal region was enriched with intercellular spaces, whereas the apical region was densely packed with dividing nuclei and apical cell processes (Figure 4A). Similar intercellular spaces have been reported in fixed specimens of developing rodent brains^44,45^. Live imaging with dextran dye confirmed the presence of basal intercellular spaces enriched in water, thereby ruling out potential artifacts from TEM sample fixation and embedding (Figure 4B).

**Figure 4.**
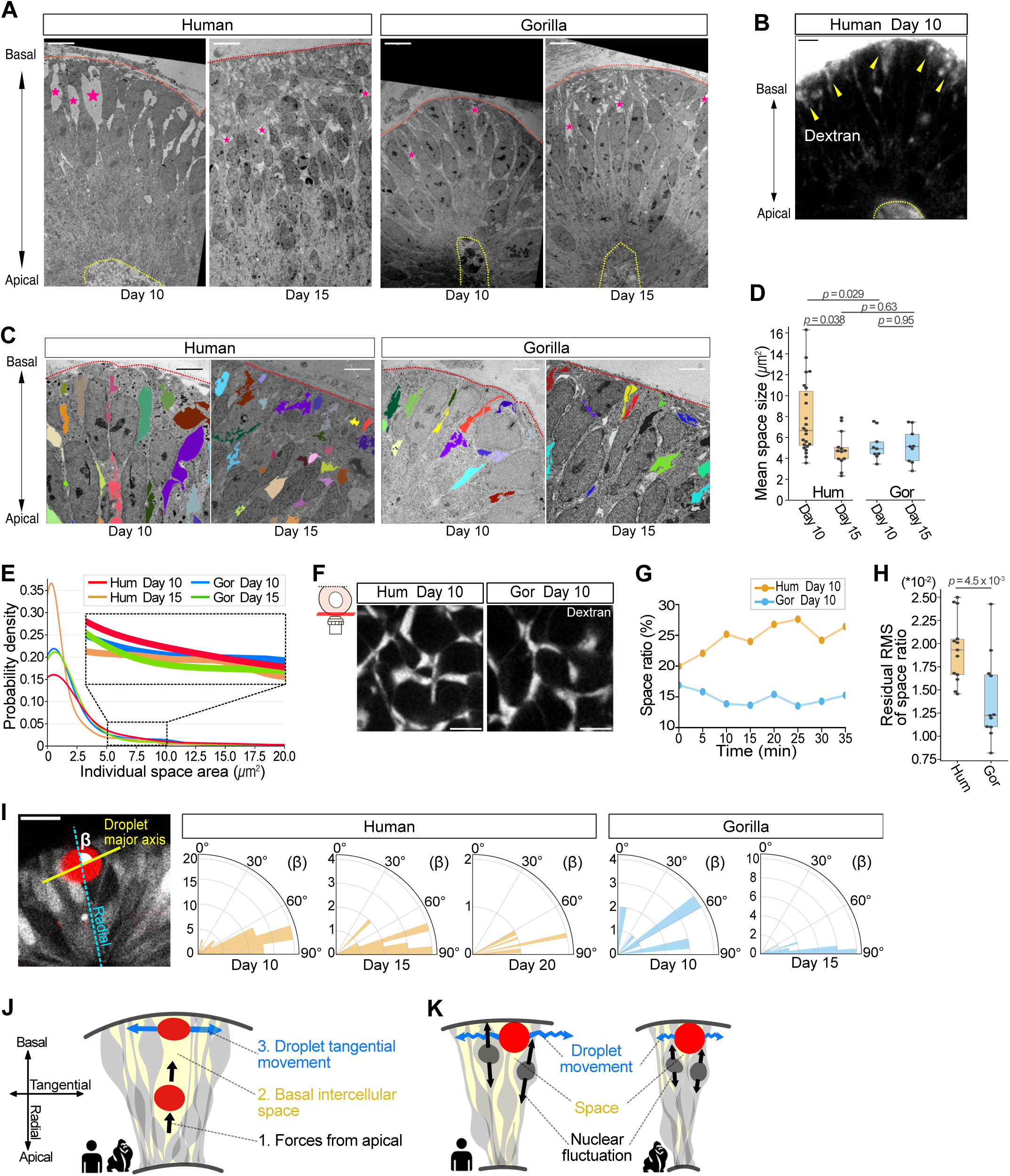
Abundant basal intercellular spaces characterize a permissive environment in human neuroepithelia (A) Transmission Electron Microscopy (TEM) images at 440× magnification of human and gorilla organoids at days 10 and 15, showing basal intercellular spaces (magenta stars). Yellow dashed outlines mark the apical surface. Scale bar: 10 μm. (B) Dextran labeling (white) of basal intercellular spaces (yellow arrowheads) in a live day 10 human organoid. Scale bar: 20 μm. (C) TEM images at 1400× magnification of human and gorilla organoids at days 10 and 15. Individual basal intercellular spaces were segmented and pseudo-colored. Scale bar: 5 μm. (D) Mean space size across species and stages. Values represent the mean size of all individual spaces per image region. human day 10 (n = 23 neural buds, N = 2 independent organoid batches), human day 15 (n = 13, N = 1), gorilla day 10 (n = 9, N = 2), gorilla day 15 (n = 9, N = 2). (E) Probability density distributions of individual space sizes (< 20 μm² shown; total range up to ∼70 μm²). Inset shows the 5-15 μm² range, where day 10 human organoids display a higher probability of containing spaces larger than 5 μm². Data as in (D). (F) Dextran labeling (white) of basal intercellular spaces in live human and gorilla organoids at day 10, imaged at the basal surface planes. Scale bar: 5 μm. (G) Representative 35-min traces of the space ratio in human and gorilla organoids. The space ratio is defined as the total basal space area divided by the entire imaged tissue area. (H) RMSE of space ratio in human and gorilla organoids. Residuals were calculated as the difference between observed and linear-regression-predicted values; higher RMSE reflects greater variability of space ratio over time, indicative of active space remodeling. Human (n = 11 tissues), gorilla (n = 11). (I) Orientation of droplet deformation angle (β) (left) and its polar plots (right) in human and gorilla organoids across stages. β is defined as the angle between the droplet major axis (yellow line) and the neural bud radial axis going through the droplet centroid (blue line). Values near 0° indicate radial alignment (i.e, forces along the tangential axis), while values near 90° indicate tangential alignment (i.e, forces along the radial axis). Polar plots include only droplets with eccentricity ≥ 0.35. Each measurement represents one droplet at one time point. Bin size is 10. Human day 10 (n = 59 droplets), human day 15 (n = 17), human day 20 (n = 7), gorilla day 10 (n = 11), gorilla day 15 (n = 33). Scale bar: 20 μm. (J) Schematic of tangential droplet movement. Droplets are displaced by apical forces toward the basal region, where intercellular spaces are available, independent of the initial injection site. Radial droplet displacement is constrained by boundaries formed by the basement membrane and extracellular matrix, thereby redirecting movement into a tangential orientation. (K) Schematic of species-specific tangential droplet movement. In human neuroepithelia, droplets exhibit longer trajectories and higher speeds, associated with greater nuclear fluctuations and larger basal spaces, whereas in gorilla neuroepithelia, droplet trajectories are shorter and speeds are lower.

In human neuroepithelia on day 10, intercellular spaces were broadly distributed in size, with several exceeding nuclear size, but these became fragmented by day 15 (Figure 4, C to E; Figure S4, A to C; see also Figure S2E). By contrast, gorilla neuroepithelia displayed smaller, more homogeneous intercellular spaces even on day 10 (Figure 4, C to E; Figure S4, A to C). Furthermore, these intercellular spaces were more dynamically remodeled over time in human neuroepithelia than in gorillas, likely stemming from greater nuclear fluctuations in humans (Figure 4, F to H; Figure S4, D to F). Together, these results suggest that larger and more dynamically remodeled intercellular spaces in the basal region may facilitate tangential droplet movement and tissue dynamics in the basal region of human neuroepithelia.

In addition to local dynamics, microdroplets are also able to quantify local anisotropic stresses within tissues^29,30^. We assessed anisotropic stresses at length scales larger than nuclear size by analyzing the ellipsoidal deformation mode of droplets^46^ (Figure S4, G and H; Methods). The microdroplets’ surface was only functionalized with PEG (Methods) to prevent cell adhesion or non-specific interactions. Consequently, droplets reported anisotropic compressive (pushing) stresses but no tensile (pulling) forces. In neuroepithelia on day 10, droplets exhibited elongated shapes corresponding to ∼0.3 kPa anisotropic stress, which is greater than the stresses previously measured in mesenchymal tissues^30^, and this anisotropic stress declined by day 15 and at later stages (Figure S4, I and J). The trend was consistent in both human and gorilla organoids. Notably, droplet elongation was oriented along the tangential axis, indicating that droplets were pushed by forces along the radial axis (Figure 4I). These radial forces, combined with the greater abundance of intercellular space in the basal region, explain the basal localization of droplets across stages and species (Figure 1I): injected droplets are pushed and displaced towards the basal region until encountering the basement membrane and Matrigel barrier, after which their movement is confined to the tangential axis (Figure 4J). This passive displacement of droplets from the apical to the basal side is consistent with the previously reported passive movement of nuclei^35^.

These data collectively reveal the mechanisms underlying droplet motility: in the basal region of human neuroepithelia, active nuclear fluctuations and permissive intercellular spaces promote tangential droplet movement, whereas at later stages or in gorilla, chimpanzee, and mouse neuroepithelia, reduced nuclear dynamics and a less permissive tissue architecture constrain droplet motility (Figure 4K). These mechanisms further suggest species-and stage-specific tissue dynamics and mechanics during forebrain development.

### Cellular rearrangements reveal basal fluidization of human neuroepithelia

While our analyses thus far have used droplet motility as a proxy for tissue dynamics and mechanics, we next studied the material state of neuroepithelia directly. We reasoned that nuclei in the basal region might also undergo tangential movement analogous to droplets, despite being more constrained by tethering to apical and basal surfaces. Indeed, basal nuclei displayed tangential movement fluctuations in addition to the previously described radial fluctuations (Figure 5A). Tangential nuclear fluctuations peaked in human neuroepithelia on day 10 and declined by day 15, whereas gorilla neuroepithelia maintained consistently low levels even on day 10 (Figure 5B). Increasing NCAD expression in human neuroepithelia suppressed tangential nuclear fluctuations (Figure 5, C and D), under which tangential droplet movement was also inhibited (Figure 3Q). This indicates that droplet motility serves as a faithful reporter of tissue dynamics. Together, these data suggest that the basal region of human neuroepithelia is more fluid than either at later developmental stages or in gorillas.

**Figure 5.**
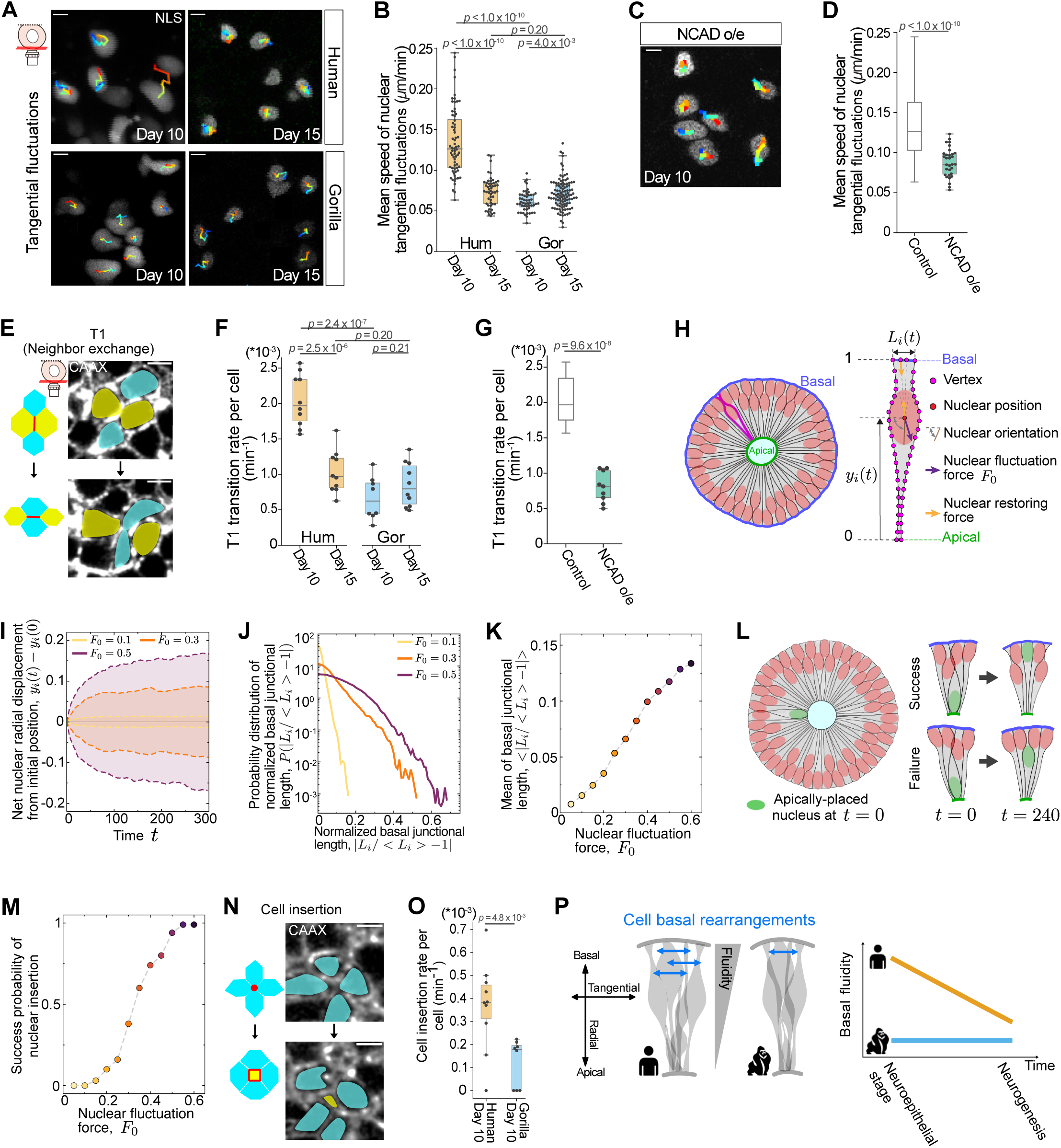
Cellular rearrangements reveal basal fluidity of human neuroepithelia (A) Representative images of basal nuclear tangential fluctuations in human and gorilla organoids at days 10 and 15, imaged at the basal surface planes. Nuclei are labeled with mCherry-NLS (white), and colored tracks show their trajectories during 3-h traces. Scale bar: 5 μm. (B) Mean speed of tangential nuclear fluctuations across species and stages. Human day 10 (n = 65 nuclei), human day 15 (n = 44), gorilla day 10 (n = 47), gorilla day 15 (n = 79). (C) Representative nuclear tangential trajectories in a day 10 NCAD o/e human organoid. Scale bar as in (A). (D) Mean speed of nuclear tangential fluctuations in NCAD o/e organoids. Control is from (B, human day 10). n = 20 nuclei. (E) Schematic of type 1 (T1) transition (left). In neighbor exchange, one cell junction shrinks while an orthogonal junction expands, resulting in a swap of neighboring cells. Representative images of T1 transitions (right), imaged at the basal surface planes. Membranes are labeled by EGFP-CAAX (white). Scale bar: 5 μm. (F) T1 transition rate per cell per min across species and stages. Human day 10 (n = 10 tissues), human day 15 (n = 10), gorilla day 10 (n = 8), gorilla day 15 (n = 10). (G) T1 transition rate in day 10 NCAD o/e organoids. Control is from (F, human day 10). n = 9 tissues. (H) Schematic of the simulation setup. The nuclear fluctuation force (*F_0_*) was modulated, and nuclear position (*y*) and the basal junctional length (*L*) were monitored. See Methods. (I) Net nuclear radial displacement at different magnitudes of nuclear fluctuation force. Dashed lines represent the standard deviation for each force condition over 10 simulations. (J) Probability distribution of normalized basal junctional length at different magnitudes of nuclear fluctuation force. (K) Mean of normalized basal junctional length at varying magnitudes of nuclear fluctuation force. (L) Schematic of the nuclear insertion simulation. A nucleus (green) was initially positioned at the apical surface (t = 0) and subsequently moved spontaneously toward the basal region. Insertion into the existing nuclear layer was classified as success if the nucleus reached the basal side at any time prior to the end of simulation. (M) Success probability of nuclear insertion at varying magnitudes of nuclear fluctuation force, calculated from 100 trials. (N) Schematic of cell insertion (left). Junctions form and expand to accommodate a new cell intercalating into the basal region. Representative images of cell insertion (right), imaged as in (E). (O) Cell insertion rate per cell per min in day 10 human and gorilla organoids. Data as in (F). (P) Schematic of species-and stage-specific basal fluidization. In early human neuroepithelia, elevated nuclear fluctuations and abundant intercellular spaces promote cell rearrangements in the basal region, including T1 transitions and cell insertion. This basal fluidity declines with the onset of neurogenesis in humans, whereas gorilla organoids exhibit consistently lower basal fluidity. Rearrangement frequencies are indicated by the number of double arrowheads.

To further assess basal fluidization, we quantified neighbor exchanges on the basal surface, or Type 1 (T1) transitions, a well-established index of tissue fluidity^47–50^ (Figure 5E). Human neuroepithelia on day 10 exhibited the highest frequency of basal T1 transitions compared with those at later developmental stages or in gorillas (Figure 5F; Figure S5, A and B; Video S5). NCAD overexpression significantly reduced basal T1 transitions (Figure 5G; Figure S5, A and B; Video S5). These findings demonstrate that the basal region of human neuroepithelia has higher cell rearrangement rates and increased tissue fluidity.

Guided by these results, we computationally studied the role of nuclear fluctuations on tissue fluidization. Specifically, we adapted the Active Foam simulation framework^51,52^ to simulate the dynamics of pseudostratified neural epithelia with nuclear dynamics while maintaining a constant cell number (Figure 5H; Methods). Increasing the magnitude of nuclear force fluctuations, F_O_, led to larger amplitudes of radial fluctuations in their positions (Figure 5I), mirroring the observed larger movement fluctuations. These larger basal nuclear fluctuations caused a greater variability in cell-cell junctional lengths on the basal side (Figure 5, J and K; Video S6), which is known to drive tissue fluidization by actively promoting T1 transitions^51^. These simulations confirm that elevated nuclear fluctuations in the basal region increase basal tissue fluidity.

We next simulated how differential tissue fluidity influences cell behavior. Because cell divisions occur exclusively at the apical surface, newly divided nuclei must relocate from the apical to basal region and be incorporated into the existing nuclear layer on the basal side. To understand the role of nuclear fluctuations in this process, we simulated the dynamics of a newborn nucleus starting on the apical side and moving towards the basal side, without explicitly implementing cell division (Figure 5L). We found that the probability of successful nuclear insertion into the basal nuclear layer increases with the magnitude of nuclear fluctuations (Figure 5, L and M; Video S7). These simulations suggest that basal nuclear fluctuations and the associated fluidization facilitate the insertion and accommodation of newborn nuclei within the basal region.

To test this prediction, we quantified cell insertion events in the basal region of neuroepithelia (Figure 5N). Human neuroepithelia on day 10 exhibited a higher rate of cell insertion compared with gorilla neuroepithelia (Figure 5O). While the measured cell insertion rate reflects both the cell division rate and the success probability of cell insertion, our simulations indicate that basal nuclear fluctuations and associated tissue fluidization increase the cell insertion rate even without changes in the division rate. These findings suggest that basal fluidization promotes tangential expansion of the basal surface in human neural buds (Figure 1D) and may ultimately contribute to human-specific forebrain expansion prior to neurogenesis.

## Discussion

In this study, we show that the basal region of early human neuroepithelia undergoes transient fluidization (Figure 5P). This basal fluidization is driven by active nuclear fluctuations and facilitated by permissive intercellular spaces. Notably, basal fluidization diminishes before neurogenic stages and appears to be specific to humans among the four species tested. The nuclear fluctuations underlying basal fluidization promote the accommodation of newly divided nuclei within the basal region and may facilitate rapid tangential expansion of the human forebrain.

Our study identifies transient basal fluidization as a previously unrecognized hallmark of early human forebrain development. While tissue fluidization has been studied in the context of tissue morphogenesis and phase transitions during embryogenesis^30,41–43,50,53–56^, it has not been systematically compared across species. Building on prior cross-species studies focused on genetic and cellular differences^2–4,21^, our findings position tissue fluidity as a biophysical parameter underlying interspecies variation in forebrain development.

Basal fluidization was revealed by the unexpected motility of oil microdroplets in cerebral organoids. The microdroplet method, originally designed to measure stress distribution^29^, serves as a simple and powerful reporter of tissue dynamics. Cerebral organoids from multiple species also provide a unified and comparable platform for mechanical analysis. Although organoid models inevitably differ from in vivo brains, they remain indispensable for cross-species comparisons, since live imaging of early human embryos (conception weeks 5-7) is challenging and not feasible for critically endangered great apes. In this context, the similarity among gorilla, chimpanzee, and mouse phenotypes is reassuring, yet expanding analyses to additional species will be necessary to further refine the phylogenetic trajectory of basal fluidization.

Our live imaging, chemical and genetic perturbations, and simulations collectively demonstrate that basal fluidization is driven by basal nuclear fluctuations. Although basal fluidization also correlates with cell division rate, increased cell division at the apical surface does not necessarily increase T1 transitions (neighbor exchanges) on the basal surface. Moreover, nuclear fluctuations are confined to the basal region, whereas nuclear movements in the apical region are predominantly directional; our simulations further show that basal nuclear fluctuations are sufficient to drive basal fluidization. NCAD overexpression suppresses both nuclear fluctuations and basal fluidization, consistent with its reported role in fluid-to-solid transitions during axis elongation in zebrafish^30^. Nevertheless, NCAD alone is unlikely to account for the phenomenon, given extensive interspecies differences in signaling and gene expression. For example, basal processes in human neuroepithelia may be more compliant and less firmly anchored, potentially permitting greater freedom for cell rearrangements. In addition, we identify abundant and permissive intercellular spaces in the basal region of human neuroepithelia, which may arise from low nuclear density or from space generated by nuclear displacement during IKNM. Since bud curvature does not significantly influence droplet motility, basal fluidization likely reflects intrinsic tissue properties rather than geometric constraints.

The discovery of basal fluidization suggests a potential mechanism underlying the rapid tangential expansion of the basal surface in the early human forebrain. Although tangential expansion has been largely attributed to the number of symmetric divisions in the apical region, and human neuroepithelia exhibit increased division rates, our findings indicate that increased nuclear fluctuations in the basal region promote the accommodation of newborn nuclei even at comparable division rates. This enhanced nuclear accommodation, together with cell divisions, can facilitate tangential surface expansion. In parallel, although the origin of basal nuclear fluctuations remains unclear, IKNM during the cell cycle may contribute to these fluctuations. Thus, basal fluidization, nuclear fluctuations, and cell division could be interconnected. While our simulations allowed independent modulation of nuclear fluctuations and basal fluidization, decoupled from cell division, achieving specific experimental control of these individual parameters remains a key challenge. Overall, our study establishes basal fluidization as a previously underappreciated dimension of evolutionary neurobiology.

## Methods

### Stem cell culture

Human IMR90-4 iPSCs (WiCell), human CRTDi011-A iPSCs^57^ (Center for Regenerative Therapies Dresden), gorilla clone 1 iPSCs^58^, chimpanzee Sandra-A iPSCs^13^, and mouse E14 ESCs^31^ were used in this study. All cells were maintained in a humidified incubator at 37 °C and 5% CO□ on plates coated with growth factor-reduced Matrigel (Corning, 356231). Cells were passaged approximately every 4 days, or when ∼80% confluent, by 3-min incubation with Accutase (Gibco, A1110501). Cells were replated at a 1:10 ratio with 10 μM ROCK inhibitor Y-27632 (StemCell Technologies, 72308) added for the first 24 h. Media were changed daily. Human, gorilla, and chimpanzee iPSCs were cultured in StemFlex™ medium (Fisher Scientific, A3349401). Mouse ESCs were initially maintained under standard 2iLIF conditions but were subsequently adapted to StemFlex medium supplemented with 1000 U/mL recombinant mLIF (Sigma-Aldrich, ESG1107) for three passages before organoid generation. All experiments involving human iPSCs were approved by the University Hospital Ethical Review Committee (IRB00001473; IORG0001076; ethical approval number SR-EK-102022025 and SR-EK-456092021).

### Cerebral organoid culture

Human and gorilla cerebral organoids were generated using identical protocols^5,11^. Reagents were prepared with the STEMdiff™ Cerebral Organoid Kit (StemCell Technologies, 08570). On day 0, iPSCs were dissociated with Accutase (5 min, 37 °C) and seeded at 500 or 2000 cells per well in ultra-low attachment U-bottom 96-well plates (Corning, 7007) in embryoid body (EB) medium containing 10 μM Y-27632. To balance handling considerations and imaging requirements, organoids generated from 2000 cells were used for immunostaining and TEM/LM analyses owing to their larger size, whereas those generated from 500 cells were used for live imaging to optimize imaging conditions. Medium was refreshed on day 3 (without ROCK inhibitor), switched to Induction Medium on day 5, and to Expansion Medium supplemented with 10% (v/v) Matrigel on day 7. Organoids were transferred to ultra-low attachment 24-well plates (Corning, 3473) with Maturation Medium on day 10. On day 14, Matrigel was removed using Cell Recovery Solution (Corning, 354253), and organoids were transferred to 5-cm dishes on orbital shakers (65 rpm). Maturation Medium was replaced every 3 days thereafter.

Chimpanzee cerebral organoids were generated following the same protocol as described above, with one modification: during the initial 2 days (day 0-day 2), cells were aggregated and cultured in mTeSR1 medium (Stem Cell Technologies, 85850) supplemented with 10 μM Y-27632. From day 3 onward, the protocol was identical to that used for human and gorilla organoids.

Mouse cerebral organoids were generated following an established protocol comparable to the human and gorilla cultures^31^. On day 0, 1000 or 2000 ESCs were seeded per well in the plates with EB medium containing 50 μM Y-27632. Medium was switched to Induction Medium on day 2 and to Expansion Medium supplemented with 10% Matrigel on day 5. On day 6, organoids were maintained in Improved Differentiation Medium without vitamin A (IDM-A)^31^. On day 7, Matrigel was removed, and organoids were transferred to orbital shakers in IDM-A. On day 8, medium was replaced with IDM formulated with B-27 supplement containing vitamin A (Gibco, 17504044) and 400 μM L-ascorbic acid (Sigma-Aldrich, A4403) and refreshed every 3 days.

### DNA constructs and reporter cell lines

The following constructs were generated for this study: pCAG-iRFP-NLS, pCAG-mCherry-NLS, pCAG-EGFP-CAAX, and pCAG-human CDH2. Coding and promoter sequences were subcloned into pDONR entry vectors and recombined into piggyBac destination vectors^59^ using Multisite Gateway cloning (Invitrogen). Constructs were stably introduced into iPSCs or ESCs by electroporation with a 4D Nucleofector (Lonza). Reporter-expressing cells were sorted either by fluorescence-activated cell sorting (FACS) or by manual clonal selection.

### Organoid epithelization labeling

To visualize neural epithelization and budding, organoids were incubated overnight with the SiR-actin probe (Spirochrome, SC001), diluted 1:1000 directly into the culture medium.

### Drug treatment

To arrest the cell cycle in S phase and inhibit nuclear movement, cerebral organoids were treated with 30 mM hydroxyurea (Sigma-Aldrich, H8627) and 150 μM aphidicolin (Sigma-Aldrich, A4487) in organoid culture medium, and incubated for 3 h. Following treatment, samples were either processed directly for live imaging (without washout) or washed before fixation for immunostaining.

### Fixation, cryosectioning, and immunostaining

Organoids were fixed in 4% paraformaldehyde (PFA; Thermo Scientific, 28906) at 4 °C overnight, washed in PBS, cryoprotected in 30% sucrose, embedded in Tissue-Tek® O.C.T.™ Compound (Sakura, 4583), and stored at-20 °C until sectioning. Sections were cut at 10 μm thickness, collected on slides and stored at-20 °C until further processing.

Cryosections were permeabilized and blocked in PBS containing Triton X-100 and donkey serum, then incubated overnight at 4 °C with primary antibodies diluted in blocking buffer. The following antibodies were used: SOX2 (R&D Systems, MAB2018-SP, 1:200), PAX6 (Abcam, ab196045, 1:200), TUJ1 (Covance, MMS-435P, 1:750), TBR1 (Abcam, ab31940, 1:300), N-Cadherin (BD Biosciences, 610920, 1:300), and Histone H3 (Abcam, ab10543, 1:300). After washing, sections were incubated with Alexa Fluor secondary antibodies (488, 568, 647; Invitrogen, 1:750) and DAPI (Invitrogen, 62247, 1:2000). Slides were mounted in ProLong™ Diamond Antifade Mountant (Thermo Fisher, P36961) for imaging. To ensure that neural buds were analyzed perpendicular to the apical-basal axis, sections displaying a clearly visible lumen and cells oriented perpendicular to the apical surface were selected.

### Generation, characterization and injection of oil microdroplets

Oil microdroplets were generated as described^29,30^. FCy5 dye^32^ was dissolved in Novec 7300 fluorocarbon oil (3M) at 0.25 mM, together with 2 wt% Krytox-PEG(600) fluorosurfactant (008-fluorosurfactant, RAN Biotechnologies). Interfacial tension was measured in cerebral Expansion Medium using pendant drop tensiometry as described^33^ using the Drop Shape Analyzer DSA25E (Krüss). The measured values were computed in a custom Python script, yielding 7.990 ± 0.627 mN/m (mean ± SD). Droplets were directly injected into organoid neural tissue with borosilicate glass capillaries (Harvard Apparatus, 30-0019). Four to six droplets were delivered per organoid. Following injection, organoids were incubated for at least 5 h to allow tissue recovery before imaging.

### Confocal imaging

Samples were imaged on Olympus FV3000 or Zeiss LSM 980 confocal microscopes. Fixed organoid sections were imaged with 10× or 20× air objectives. Live cerebral organoids were imaged in glass-bottom chambers with either a 30× silicone oil objective (UPLSAPO30X, Olympus FV3000) or a 40× water-immersion objective (LD C-Apochromat 40×/1.1, Zeiss LSM 980) inside an incubator (37 °C, 5% CO□). For droplet deformation imaging, the Zeiss LSM 980 was used in Airyscan 2 mode (32-channel GaAsP detector) at optimal Z-resolution for droplet 3D reconstruction, while standard confocal mode was used for droplet tracking.

For images of the basal tissue region, z-stacks were acquired at the organoid surface planes. This included dextran imaging of basal intercellular spaces, membrane imaging of cell rearrangements, and nuclear tracking for their tangential fluctuations. For images showing tissue apical-basal axis, z-stacks were acquired around the lumen, with focus planes centered on the detected lumen planes, and imaging planes selected following the criteria described in *Fixation, cryosectioning, and immunostaining*. Time-lapse imaging intervals were 5 min for droplet dynamics and intercellular space dextran-labeling, and 10 min for nuclear movements and cell rearrangements (membrane labeling). Images were processed in Fiji (ImageJ) or with custom Python scripts.

### Electron and light microscopy

Cerebral organoids were fixed in 1% glutaraldehyde, 2% paraformaldehyde, and 0.08 mM CaCl□ in 0.1 M phosphate buffer (pH 7.4) for 1 h at room temperature, followed by overnight fixation at 4 °C. Samples were washed three times in 0.08 mM CaCl□ in phosphate buffer and once in ddH□O, stained in 2% osmium tetroxide with 1.5% potassium ferricyanide for 1 h at room temperature, washed in ddH□O, dehydrated through a graded ethanol series, and embedded in Epon 812 resin (Electron Microscopy Sciences).

From this point, samples were processed for either light microscopy or transmission electron microscopy (TEM). For light microscopy, 1.5-μm sections were sliced, stained with toluidine blue, and imaged on an Axio Imager.Z2 microscope (Zeiss) with a 63× air objective. For TEM, ultrathin sections (∼70 nm) were collected on Formvar-coated grids, post-stained with uranyl acetate and lead citrate, and imaged on a Tecnai 12 transmission electron microscope (Thermo Fisher) operated at 100 kV and equipped with an F416 CMOS camera (TVIPS) at magnifications of 440×, 1200×, and 1400×.

### Neural bud perimeter quantification

Neural bud boundaries were manually annotated based on cell membrane, and measured in Fiji (ImageJ) using the ‘Segmented Line’ tool.

### Neural bud basal surface curvature analysis

The local basal contour of neural buds at droplet-contact regions was manually traced in Fiji using the ‘Segmented Line’ tool, followed by ‘Fit Spline’ tool to smooth the contour. Traced region-of-interest (ROIs) were exported as XY coordinates and analyzed in a custom Python pipeline. Briefly, circles were fitted to the local contours using least-squares minimization. The local radius of curvature (R) was defined as the mean fitted radius, and curvature was calculated as its reciprocal (1/R). For each ROI, fitted contours and circle overlays were plotted for manual inspection, and summary statistics (radius, curvature) were saved to Excel.

To test functional relevance between droplet motility and tissue surface curvature, droplet motility was paired with local curvature. ROI-derived curvature values were merged with droplet motility data, and Pearson’s correlation coefficients (r, p) were computed.

### Ellipsoidal stress quantification

Ellipsoidal stresses were quantified from droplet deformations using the STRESS software package^46^. Briefly, 3D droplet shapes were reconstructed from confocal image stacks, and anisotropic stresses (σ_T_^A^) were computed from the ellipsoidal mode of deformation via a least-squares ellipsoidal fit:

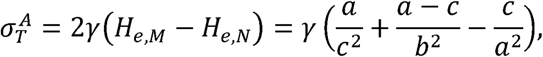

where H_e,M_ and H_e,N_ are the maximum and minimum mean curvatures of the ellipsoid, and r is the droplet interfacial tension; and a, b, and c are the major, medial, and minor axes of the ellipsoid. Reconstructed droplet was visualized in Paraview.

### Droplet tracking and motion analysis

Sample drift was corrected with the MultiStackReg plugin in Fiji^60^, using the fluorescent membrane channel as a reference to align droplet images across frames. Droplet trajectories were then extracted with the TrackMate plugin^61^ using Laplacian of Gaussian detection. Only trajectories spanning ≥ 150 min were included in the analysis. TrackMate outputs included track mean speed, displacement per interval, instantaneous speed, and centroid coordinates for each droplet at every time point.

### Analyses of droplet size, position, and angles of deformation and movement

Droplet and tissue features were quantified from confocal images using a custom Python script. Droplets were segmented by intensity thresholding followed by contour detection, and the largest connected component was selected to isolate the droplet. Droplet size was calculated from the cross-sectional area at the manually determined central plane. The droplet radius was calculated as the mean of the fitted ellipse axes:

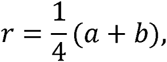

where a and b are the major and minor axis lengths, respectively.

For positional analysis, two reference points were manually annotated along the radial axis passing through the droplet centroid: P1, at the basal boundary of the ventricular zone (VZ), and P2, at the apical surface (lumen). VZ tissue thickness was defined as the distance between P1 and P2. Droplet position was measured as the distance from the droplet centroid to the apical surface (P2) and normalized to the VZ thickness (0 = apical, 1 = basal).

The droplet deformation angle (β) was defined as the angle between the major axis of the fitted ellipse and the tissue’s radial axis passing through the droplet centroid. Angles near 0° correspond to radial alignment (forces acting tangentially), whereas angles near 90° correspond to tangential alignment (forces acting radially). For each condition (stage × species), 9 droplets were analyzed over the first 11 time points. Only droplets with an eccentricity ≥ 0.35 (i.e., with detectable deformation) were included in this analysis.

Droplet movement direction was analyzed by tracking centroid positions across consecutive frames. Displacement vectors were computed between centroids at T and T + 4. The local radial axis was defined as the line connecting P1 and P2 at T. Movement angle (θ) was quantified as the angle between the displacement vector and this radial axis, and constrained to 0°-90°, with 0° indicating radial movement and 90° tangential movement. Angles were only calculated when (i) ≥ 4 frames of trajectory history were available, and (ii) droplet displacement exceeded half the droplet diameter.

### Nuclear tracking and motion analysis along the tangential axis

Tangential nuclear movement was analyzed using the same pipeline as described above for droplet tracking. Only trajectories lasting ≥ 100 min were retained. Track mean speed was extracted for analysis.

### Nuclear tracking and motion analysis along the radial axis

Sample drift was corrected, as described above for droplet tracking. Nuclear trajectories were determined using the Manual Track plugin. For nuclear radial fluctuations, only trajectories lasting at least 150 min were retained, and the first 150 min of each trajectory were analyzed. For nuclear migrations from the apical surface, only trajectories lasting at least 40 min were retained, and analysis was restricted to the first 40 min, right after mitosis with nuclei still within 30 μm of the apical surface. From these trajectories, displacement per interval, instantaneous speed, and centroid coordinates were extracted. Nuclear mean speed was calculated as the sum of all interval displacements divided by the total track duration.

Net displacement from the initial position was calculated by setting the droplet position at t=0 as the reference point. Displacement vectors were defined as 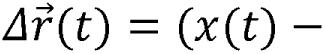 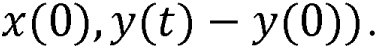 For plotting, displacements were projected onto the radial axis, allowing positive and negative values relative to the initial position. Net displacement traces were fit with linear regression, and the root-mean-square error (RMSE) of the residuals was calculated from the residuals between observed and predicted values.

### Segmentation and quantification of intercellular space in TEM images

Intercellular space was analyzed using a custom Python script. Segmentation was challenging due to (i) markedly heterogeneous intensity across the field, (ii) highly irregular and diverse morphologies, and (iii) frequent small noise artifacts. Under these conditions, conventional algorithms such as global intensity thresholding (e.g., Otsu), watershed segmentation, and gradient-or edge-based methods perform unreliably. Deep learning approaches likewise fail to generalize. Given that local intensity is relatively uniform within small regions, a sliding-window adaptive local-thresholding algorithm was implemented. Each image was divided into overlapping windows, and an intensity histogram was computed for each window following contrast equalization. For windows displaying multimodal histograms, Otsu’s threshold was applied with a conservative offset to avoid over-segmentation of faint structures; whereas for histograms lacking bimodality, an empirically chosen percentile-based threshold was applied. To limit deviations caused by noise or intensity spikes, local thresholds were constrained by comparison to a global percentile threshold estimated from the full image. Afterwards, binary masks from overlapping windows were combined by union to produce the final segmentation, preserving fine-grained local adaptivity while ensuring spatial continuity across the full image. Segmentation masks were exported and visualized in Napari, and then manually inspected and corrected using a custom plugin to ensure accuracy in downstream quantitative analyses.

For each image, the script returned individual intercellular space sizes, mean space size, and total space count. Segmented spaces smaller than 20,000 pixels (≈1.53 μm²) were excluded. For statistical analysis, probability density functions were estimated from pooled distributions of individual space sizes within each condition (stage x species).

### Intercellular space labeling and segmentation with dextran

To visualize intercellular spaces in live cerebral organoids, Dextran-Alexa Fluor 647 (Invitrogen, D22914) was added to the culture medium at 5 μM. After 1.5-h incubation, confocal time-lapse imaging was performed. Regions of 20.09 × 20.09 μm² were cropped for analysis, via a custom Napari plugin.

Considering the high signal-to-background contrast and clean intensity distributions, global thresholding provided reliable segmentation. Otsu’s method with an empirically defined offset was then applied frame by frame to generate binary masks. Masks were overlaid with the raw images in Napari for inspection, and segmentation errors (e.g., incomplete thresholding or small artifacts) were corrected manually using Napari’s interactive labeling tools. Final masks were exported for downstream quantification. Space ratio was defined as the fraction of image area occupied by segmented intercellular space (i.e., total space area / total image area). Temporal variability was quantified by fitting the space ratio time series with linear regression and computing RMSE of the residuals.

### Cell division rate analysis

Mitotic figures were identified in light microscopy images based on their apical localization and characteristic chromatin condensation. The number of dividing cells was normalized to the lumen perimeter, calculated as the count of mitotic figures divided by the lumen perimeter length.

### N-cadherin fluorescence intensity profiling

Confocal Z-stacks were acquired with the same parameters and projected by ‘Sum Slices’. Fluorescence intensity profiles of N-cadherin were obtained by drawing a line, with a width of 10 pixels, from the apical to the basal surface in Fiji (ImageJ) and generating the corresponding plot profile of gray values. Line lengths were normalized from 0 to 1, where 0 represents the apical surface and 1 the basal surface.

### Cell rearrangements analysis

Cell rearrangement events refer to junctional changes in altered neighboring relationships, and were manually identified and classified in time-lapse membrane imaging. Live imaging was performed for at least 100 min. Regions of 40.10 × 40.10 μm² were cropped for analysis. Rearrangement rates were normalized to both the total number of cells in the analyzed region at t= 0 and the imaging duration. Cumulative rearrangements over time were quantified by summing events up to each time point and normalizing to the initial cell number.

### Simulations of neural epithelia tissue and nuclear dynamics

Based on the previously developed Active Foam Model with nucleus^51,52^, we extend the model by modifying the geometric description and cell-cell and cell-nucleus mechanical interactions to investigate organoid systems. Cell junctions are classified into apical, lateral, and basal sides. The corresponding junctional tensions are defined such that the apical and basal tensions are proportional to the respective junctional lengths, with distinct moduli K_T,a_ and K_T,b_, while the lateral tension is assumed to be constant, T_l_= T_O_. The tensions are given by,

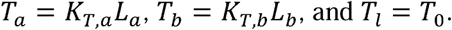

The cell aspect ratio is determined by the energetic balance between apical, lateral, and basal tensions. An increase in apical and basal tensions relative to the lateral tension leads to a higher aspect ratio. K_T,a_, K_T,b_, and T_O_ are selected to yield a high cell aspect ratio consistent with pseudostratified epithelial structure observed experimentally. The inner lumen is modeled as a cell with a distinct osmotic pressure.

To incorporate interkinetic nuclear dynamics, we introduce two effective forces on the nucleus, namely a nuclear restoring force and a nuclear fluctuation force. The nuclear restoring force represents the interaction between the nucleus and the basal junction, promoting basal positioning of the nucleus. This force is implemented as a spring-like elastic interaction between the nuclear center and each vertex on the basal junction of a given cell,

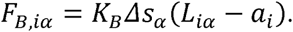

Here, K_B_ denotes the restoring force modulus, Lls_a_ is the normalized segment length associated with vertex a on the basal junction, L_ia_ is the distance between the nucleus of cell i and vertex a on its basal junction, and a_i_ is the length of semi-major axis of a given nucleus i.

The nuclear fluctuation force is modeled as an active Brownian particle characterized by a magnitude F_O_ and a persistent timescale, T_p_. The competition between the restoring and fluctuation forces results in deviations of the nuclear position away from the basal side.

All forces are non-dimensionalized using characteristic scales as described in previous work^51,52^, and the system is numerically integrated using the Euler-Maruyama method. For all simulations, the initial configuration is generated as an annulus geometry with prescribed inner and outer radii. This configuration is first relaxed in the absence of nuclear fluctuation forces for a duration of t= 20. Following this initial relaxation, a finite nuclear fluctuation force is applied, and the system is further evolved for an additional t= 20 before data sampling.

For simulations involving an apically positioned nucleus, one randomly selected nucleus is constrained to the apical side during the initial sampling period of t= 20, then the constraint is removed. All simulations are performed for a total duration of t= 400.

## Statistical analysis

All biological experiments were performed using three independent organoid batches, unless otherwise specified in the figure legends. Sample sizes used are reported in the respective legends.

Normality was assessed with the Shapiro-Wilk test. If both groups followed a normal distribution, equality of variances was tested with Levene’s test. For equal variances (*p* > 0.05), a two-sided Student’s *t*-test was applied; for unequal variances (*p* ≤ 0.05), a two-sided Welch’s *t*-test was used. If at least one group deviated from normality, a two-sided Mann-Whitney U test was performed as a non-parametric alternative. For multiple comparisons, p-values were adjusted using the Bonferroni correction. All analyses were conducted in Python.

## Acknowledgments

This work was supported by EMBL; the Deutsche Forschungsgemeinschaft (DFG, German Research Foundation) under Germany’s Excellence Strategy - EXC 2068 - 390729961 - Cluster of Excellence Physics of Life of TU Dresden; and the European Research Council (ERC) under the European Union’s Horizon 2020 research and innovation program (grant agreement no. 101002564, to M.E.). M.E. is supported by the Alexander von Humboldt Foundation through the Alexander von Humboldt Professorship endowed by the Federal Ministry of Education and Research. S.X. is supported by the Dresden International Graduate School for Interdisciplinary Life Sciences (DIGS-ILS).

The human CRTDi011-A iPSC line^57^ and its parental fibroblasts were provided by the Stem Cell Engineering Facility at the Center for Regenerative Therapies Dresden (CRTD) and by Nataliya Di Donato, respectively. The gorilla iPSC line^58^, chimpanzee iPSC line^13^, and mouse ESC line^31^ were kindly provided by Ulrich Martin, Svante Pääbo, and Madeline Lancaster, respectively.

We would like to thank the following Services and Facilities for their technical support: MPI-CBG Organoid and Stem Cell Facility, MPI-CBG Electron Microscopy, MPI-CBG Light Microscopy, EMBL Barcelona Mesoscopic Imaging Facility, PoL Microscopy Facility, CRG Tissue Engineering Unit, and CMCB Histology Facility.

We thank Ellen Sletten for sharing custom-made FCy5 dyes; Madeline Lancaster for sharing organoid protocols; Mitsuhiro Matsuda for generating plasmids and supervising cell and organoid culture; Guillermo Martínez-Ara for supervising cell and organoid cultures; Ziang Ma for designing animal icons; Manon Valet for advising on cell rearrangement analysis; Claudia Fröb for generating oil microdroplets; Shuo-Ting Yen, Marc Trani Bustos, and Johannes Soltwedel for guiding droplet analysis; Ebisuya and Trivedi laboratories for support; Caren Norden for manuscript feedback.

## Author contributions

S.X. and M.E. designed the work and wrote the manuscript. S.X. performed most experiments and analyzed all data. S.K. performed the simulations. G.L. developed pipelines for space segmentation and quantification. M.W.-B. advised on TEM preparation and interpretation. A.K., T.M.S., and M.A. contributed to chimpanzee and human (CRTD) cerebral organoids. O.C. provided oil microdroplets, guided the mechanical interpretation of the data, and supervised the analysis and simulation of tissue fluidity. V.T. and M.E. supervised the project. All authors contributed to the manuscript and approved the final version.

## Generative AI and AI-assisted technologies

AI tools were used to improve text clarity and assist with code generation. The authors reviewed and edited all outputs and assume full responsibility for the content.

## Lead contact

Requests for further information and resources should be directed to and will be fulfilled by the lead contacts, Miki Ebisuya or Otger Campàs.

## Materials availability

All cell lines were previously published or available from commercial sources. Other materials are available from the lead contact.

## Data and code availability

All custom code developed and used in this study is available at: https://github.com/mebisuya/BrainFluidity https://github.com/MESOBIO-EPFL/2026_ORGANOID_NUCLEAR_DYNAMICS

## Supplementary figure legends

Figure S1. Human neuroepithelia display high droplet instantaneous speed, related to Figure 1

(A) Bright-field images of human, gorilla, and mouse pluripotent stem cells. ESCs, embryonic stem cells; iPSCs, induced pluripotent stem cells. Scale bar: 100 μm.

(B) SiR-actin labeling of neural bud morphology transformations during initial stages of organoid epithelization. Neural buds are incomplete and not yet well segregated at the indicated stages. Scale bar: 50 μm.

(C) Droplet radius across species and stages. Each data point represents one droplet measured at one time point. Data as in Figure 1I.

(D) Droplet instantaneous speed traces of all analyzed droplets across species and stages (left), and corresponding mean ± SD (right). Each line represents one droplet trace. Red arrowheads highlight large-displacement events (“jumps”). Data as in Figure 1H. Human IMR90 day 10 (n = 17 droplets), human day 15 (n = 49), human day 20 (n = 32), human CRTD day 10 (n = 34), gorilla day 8 (n = 18), gorilla day 10 (n = 12), gorilla day 15 (n = 20), chimpanzee day 10 (n = 9), mouse days 6/6.5 (n = 17).

(E) Droplets in day 8 gorilla cerebral organoids (top) with corresponding 3-h color-coded trajectories (bottom). Membranes are labeled by EGFP-CAAX (white), and the droplet is shown in red. Left, a young bud formed during initial neuroepithelialization; right, a region still undergoing epithelization thereby no bud clearly detected. Scale bar: 20 μm (top), 10 μm (bottom).

(F) Bright-field images of cerebral organoids derived from human CRTD (top) and chimpanzee (bottom) cells during neuroepithelialization. Scale bar: 100 μm.

(G) Tissue thickness across species and developmental stages. In day 20 human organoids, this corresponds to the thickness of the ventricular zone, excluding the neural layer. Data as in Figure 1I.

(H) Representative images of droplets (red) at low-and high-curvature regions of day 10 human neural buds. Membranes are labeled with EGFP-CAAX (green). Scale bar: 10 μm.

**Figure S2. Nuclei show greater radial fluctuations in human basal regions than in gorilla, related to Figures 2 and 4**

(A) Real-time displacement traces of radial fluctuations in all nuclei analyzed across species and stages. Displacement is plotted relative to the initial position at t = 0, as in Figure 2B. Each line represents one nuclear trace. Data as in Figure 2B (right).

(B) Nuclear instantaneous speed traces of all analyzed nuclei across species and stages. Red arrowheads highlight greater nuclear fluctuations observed in day 10 human organoids. Data as in (A).

(C) Real-time displacement traces of migration from the apical surface in all nuclei analyzed across species and stages (top), with mean ± SD (bottom). Data as in Figure 2F (migration from apical).

(D) Mean speed of nuclear migration from the apical surface across species and stages. Data as in (C).

(E) Light microscopy images of the basal region of human and gorilla organoids at days 10 and 15. Scale bar: 5 μm.

**Figure S3. Droplet motility is reduced under nuclear fluctuation inhibition, related to** Figure 3

(**A–B**) Real-time displacement traces of radial fluctuations in all nuclei analyzed in Hu-Aph-treated day 10 human organoids (A), with mean ± SD (B). Data as in Figure 3C. (**C–D**) Droplet instantaneous speed traces of all droplets analyzed in Hu-Aph-treated day 10 human organoids (C), with mean ± SD (D). Data as in Figure 3F.

(**E**) NCAD intensity profiles of all human, gorilla, and NCAD o/e organoids at day 10. Data as in Figure 3H.

(**F–G**) Real-time displacement traces of radial fluctuations in all nuclei analyzed NCAD o/e organoids at day 10 (F), with mean ± SD (G). Data as in Figure 3L.

(**H–I**) Droplet instantaneous speed traces of all droplets analyzed in NCAD o/e organoids at day 10 (H), with mean ± SD (I). Data as in Figure 3Q.

**Figure S4. Basal space is larger and more dynamically remodeled in early human neuroepithelia, related to** Figure 4

(A) Number of basal intercellular spaces per image region across species and stages. Data as in Figure 4D.

(B) Distribution of individual basal intercellular space sizes across species and stages. Each data point represents one space from one image region. Human day 10 (n = 858 individual spaces), human day 15 (n = 985), gorilla day 10 (n = 256), gorilla day 15 (n = 424).

(C) Probability of observing basal intercellular spaces above defined size thresholds (>5, >10, >20 μm²) across species and stages, calculated based on Figure 4E.

(D) Dextran labeling (white) of basal intercellular spaces in day 10 human and gorilla organoids, imaged at basal surface planes and shown with orthogonal sections. Scale bar: 10 μm.

(E) Real-time traces of basal space ratio in all image regions analyzed in day 10 human and gorilla organoids. Each line represents one image region trace. Data as in Figure 4H.

(F) Mean basal space ratio in day 10 human and gorilla organoids, calculated by averaging space ratio values over each trace. Data as in (E).

(G) Droplet interfacial tension. Each data point is from an independent measurement. Mean ± SD: 7.990 ± 0.627 mN/m.

(H) Schematic of the oil droplet assay for stress measurement. In suspension, droplets remain spherical; when inserted into tissue, anisotropic forces (black arrows) derived from neighboring cells deform droplets into ellipsoids^29^.

(I) 3D reconstructed droplets and mean curvature maps of ellipsoidal mode in human and gorilla organoids. Curvature was calculated using the Stress software^46^ and droplets were reconstructed in PARAVIEW. Blue indicates compression and red indicates tension, with brightness reflecting curvature magnitude.

(J) Quantification of ellipsoidal stress using the droplet assay. Each data point represents stress measured from one droplet at one time point. Human day 10 (n = 16 droplets), human day 15 (n = 28), human day 20 (n = 14), gorilla day 10 (n = 14), gorilla day 15 (n = 10).

**Figure S5. Basal cell rearrangements are pronounced in human neuroepithelia, related to** Figure 5

(**A–B**) Cumulative T1 transition counts per cell over time. Data as in Figure 5, F and G.

(A) Individual traces in human, gorilla, and NCAD o/e organoids at the indicated days with each line representing one image region. (**B**) Mean values with linear regression fits.

## Supplementary videos

**Video S1. Droplet moves along the tangential axis, related to** Figure 1

Time-lapse imaging of an injected oil droplet (red) in a day 10 human neuroepithelium. Its trajectory is color-coded, with blue marking the start and red marking the end. Nuclei are labeled with mCherry-NLS (white). Scale bar: 20 μm.

**Video S2. Droplets exhibit dynamic motility exclusively in human neuroepithelia, related to** Figure 1

Time-lapse imaging of droplets (red) in the basal region of human IMR90, human CRTD, gorilla, mouse, and chimpanzee organoids. Membranes are labeled with EGFP-CAAX (white). Scale bar = 20 μm.

**Video S3. Nuclei exhibit pronounced radial fluctuations exclusively in human basal neuroepithelia, related to** Figure 2

Time-lapse imaging of nuclei (magenta) in the basal region of human, gorilla and mouse organoids, with their color-coded trajectories. Membranes are sparsely labeled with EGFP-CAAX (white). Scale bar = 20 μm.

**Video S4. Suppression of nuclear fluctuations reduces droplet motility, related to** Figure 3

Time-lapse imaging of radial nuclear fluctuations (top panels) and corresponding droplet dynamics (bottom panels) in Hu-Aph-treated or NCAD o/e organoids. Membranes are labeled with EGFP-CAAX (green), nuclei with mCherry-NLS (magenta), and droplets in red. Hu-Aph, hydroxyurea-aphidicolin; NCAD o/e, N-cadherin overexpression. Scale bar = 10 μm.

**Video S5. Human basal neuroepithelial surface undergoes active cell rearrangements, related to** Figure 5

Time-lapse imaging of T1 transitions (green dots) in the basal region of neuroepithelia. Membranes are visualized with the SiR-actin probe (magenta). hum, human; gor, gorilla; d, day. Scale bar = 5 μm.

**Video S6. Simulated effects of nuclear fluctuations on basal junctional lengths, related to** Figure 5.

The magnitude of the nuclear fluctuation force was varied from low (F_O_ = 0.1, left), medium (F_O_ = 0.3, center), and high (F_O_ = 0.5, right), and the corresponding fluctuations in basal junctional lengths were simulated.

**Video S7. Simulated effects of nuclear fluctuations on nuclear insertion into the basal region, related to** Figure 5.

A nucleus (green) was initially constrained at the apical surface, and the constraint was removed at t= 20. The nucleus subsequently migrated toward the basal region. The magnitude of the nuclear fluctuation force was varied from low (F_O_ = 0.1), medium (F_O_ = 0.3), and high (F_O_ = 0.5), and the corresponding nuclear insertion was simulated. “Failure” and “Success” indicate whether the nucleus was successfully inserted into the existing nuclear layer in the basal region.

## References

1. Herculano-Houzel, S. & Kaas, J. H. Gorilla and Orangutan Brains Conform to the Primate Cellular Scaling Rules: Implications for Human Evolution. Brain. Behav. Evol. 77, 33–44 (2011).

2. Florio, M., Borrell, V. & Huttner, W. B. Human-specific genomic signatures of neocortical expansion. Curr. Opin. Neurobiol. 42, 33–44 (2017).

3. Vanderhaeghen, P. & Polleux, F. Developmental mechanisms underlying the evolution of human cortical circuits. Nat. Rev. Neurosci. 24, 213–232 (2023).

4. Lindhout, F. W., Krienen, F. M., Pollard, K. S. & Lancaster, M. A. A molecular and cellular perspective on human brain evolution and tempo. Nature 630, 596–608 (2024).

5. Benito-Kwiecinski, S. et al. An early cell shape transition drives evolutionary expansion of the human forebrain. Cell 184, 2084–2102.e19 (2021).

6. Rakic, P. Evolution of the neocortex: a perspective from developmental biology. Nat. Rev. Neurosci. 10, 724–735 (2009).

7. Götz, M. & Huttner, W. B. The cell biology of neurogenesis. Nat. Rev. Mol. Cell Biol. 6, 777–788 (2005).

8. Uzquiano, A. & Arlotta, P. Brain organoids: the quest to decipher human-specific features of brain development. Curr. Opin. Genet. Dev. 75, 101955 (2022).

9. Paşca, S. P. Assembling human brain organoids. Science 363, 126–127 (2019).

10. Sasai, Y. Cytosystems dynamics in self-organization of tissue architecture. Nature 493, 318–326 (2013).

11. Lancaster, M. A. et al. Cerebral organoids model human brain development and microcephaly. Nature 501, 373–379 (2013).

12. Otani, T., Marchetto, M. C., Gage, F. H., Simons, B. D. & Livesey, F. J. 2D and 3D Stem Cell Models of Primate Cortical Development Identify Species-Specific Differences in Progenitor Behavior Contributing to Brain Size. Cell Stem Cell 18, 467–480 (2016).

13. Mora-Bermúdez, F. et al. Differences and similarities between human and chimpanzee neural progenitors during cerebral cortex development. eLife 5, e18683 (2016).

14. Kanton, S. et al. Organoid single-cell genomic atlas uncovers human-specific features of brain development. Nature 574, 418–422 (2019).

15. Pollen, A. A. et al. Establishing Cerebral Organoids as Models of Human-Specific Brain Evolution. Cell 176, 743–756.e17 (2019).

16. Agoglia, R. M. et al. Primate cell fusion disentangles gene regulatory divergence in neurodevelopment. Nature 592, 421–427 (2021).

17. Cubillos, P. et al. The growth factor EPIREGULIN promotes basal progenitor cell proliferation in the developing neocortex. EMBO J. 43, 1388–1419 (2024).

18. Nolbrant, S. et al. Interspecies Organoids Reveal Human-Specific Molecular Features of Dopaminergic Neuron Development and Vulnerability. Preprint at 10.1101/2024.11.14.623592 (2024).

19. Wu, Q. et al. Epigenetic–metabolic axis in the temporal scaling of mammalian cortical neurogenesis across species. Preprint at 10.1101/2025.09.23.677958 (2025).

20. Reilly, S. K. et al. Evolutionary changes in promoter and enhancer activity during human corticogenesis. Science 347, 1155–1159 (2015).

21. Pollen, A. A., Kilik, U., Lowe, C. B. & Camp, J. G. Human-specific genetics: new tools to explore the molecular and cellular basis of human evolution. Nat. Rev. Genet. 24, 687–711 (2023).

22. Pathak, M. M. et al. Stretch-activated ion channel Piezo1 directs lineage choice in human neural stem cells. Proc. Natl. Acad. Sci. 111, 16148–16153 (2014).

23. Tallinen, T. et al. On the growth and form of cortical convolutions. Nat. Phys. 12, 588–593 (2016).

24. Karzbrun, E., Kshirsagar, A., Cohen, S. R., Hanna, J. H. & Reiner, O. Human brain organoids on a chip reveal the physics of folding. Nat. Phys. 14, 515–522 (2018).

25. Campàs, O. A toolbox to explore the mechanics of living embryonic tissues. Semin. Cell Dev. Biol. 55, 119–130 (2016).

26. Sugimura, K., Lenne, P.-F. & Graner, F. Measuring forces and stresses in situ in living tissues. Development 143, 186–196 (2016).

27. Roca-Cusachs, P., Conte, V. & Trepat, X. Quantifying forces in cell biology. Nat. Cell Biol. 19, 742–751 (2017).

28. Prevedel, R., Diz-Muñoz, A., Ruocco, G. & Antonacci, G. Brillouin microscopy: an emerging tool for mechanobiology. Nat. Methods 16, 969–977 (2019).

29. Campàs, O. et al. Quantifying cell-generated mechanical forces within living embryonic tissues. Nat. Methods 11, 183–189 (2014).

30. Mongera, A. et al. A fluid-to-solid jamming transition underlies vertebrate body axis elongation. Nature 561, 401–405 (2018).

31. Lloyd-Davies Sánchez, D. J., Lindhout, F. W., Anderson, A. J., Pellegrini, L. & Lancaster, M. A. Mouse brain organoids model in vivo neurodevelopment and function and capture differences to human. Preprint at 10.1101/2024.12.21.629881 (2024).

32. Lim, I. et al. Fluorous Soluble Cyanine Dyes for Visualizing Perfluorocarbons in Living Systems. J. Am. Chem. Soc. 142, 16072–16081 (2020).

33. Van De Wouw, H. L. et al. NonLIonic Fluorosurfactants for DropletLBased in vivo Applications. Angew. Chem. Int. Ed. 63, e202404956 (2024).

34. Norden, C., Young, S., Link, B. A. & Harris, W. A. Actomyosin Is the Main Driver of Interkinetic Nuclear Migration in the Retina. Cell 138, 1195–1208 (2009).

35. Kosodo, Y. et al. Regulation of interkinetic nuclear migration by cell cycle-coupled active and passive mechanisms in the developing brain: Mechanisms of interkinetic nuclear migration. EMBO J. 30, 1690–1704 (2011).

36. Murciano, A., Zamora, J., Lopezsanchez, J. & Frade, J. Interkinetic Nuclear Movement May Provide Spatial Clues to the Regulation of Neurogenesis. Mol. Cell. Neurosci. 21, 285–300 (2002).

37. Maia-Gil, M. et al. Nuclear deformability facilitates apical nuclear migration in the developing zebrafish retina. Curr. Biol. 34, 5429–5443.e8 (2024).

38. Campàs, O., Noordstra, I. & Yap, A. S. Adherens junctions as molecular regulators of emergent tissue mechanics. Nat. Rev. Mol. Cell Biol. 25, 252–269 (2024).

39. Hatta, K. & Takeichi, M. Expression of N-cadherin adhesion molecules associated with early morphogenetic events in chick development. Nature 320, 447–449 (1986).

40. Dady, A., Blavet, C. & Duband, J. Timing and kinetics of EL to NLcadherin switch during neurulation in the avian embryo. Dev. Dyn. 241, 1333–1349 (2012).

41. Petridou, N. I., Grigolon, S., Salbreux, G., Hannezo, E. & Heisenberg, C.-P. Fluidization-mediated tissue spreading by mitotic cell rounding and non-canonical Wnt signalling. Nat. Cell Biol. 21, 169–178 (2019).

42. Petridou, N. I., Corominas-Murtra, B., Heisenberg, C.-P. & Hannezo, E. Rigidity percolation uncovers a structural basis for embryonic tissue phase transitions. Cell 184, 1914–1928.e19 (2021).

43. Manning, M. L. Rigidity in mechanical biological networks. Curr. Biol. 34, R1024–R1030 (2024).

44. Hinds, J. W. & Ruffett, T. L. Cell proliferation in the neural tube: An electron microscopic and Golgi analysis in the mouse cerebral vesicle. Z. Zellforsch. Mikrosk. Anat. 115, 226–264 (1971).

45. Sumi, S. M. The extracellular space in the developing rat brain: Its variation with changes in osmolarity of the fixative, method of fixation and maturation. J. Ultrastruct. Res. 29, 398–415 (1969).

46. Gross, B. J., Soltwedel, J. R., Shelton, E., Gomez, C. & Campàs, O. STRESS, an automated geometrical characterization of deformable particles for in vivo measurements of cell and tissue mechanical stresses. Sci. Rep. 15, 28599 (2025).

47. Guillot, C. & Lecuit, T. Mechanics of Epithelial Tissue Homeostasis and Morphogenesis. Science 340, 1185–1189 (2013).

48. Fletcher, A. G., Osterfield, M., Baker, R. E. & Shvartsman, S. Y. Vertex Models of Epithelial Morphogenesis. Biophys. J. 106, 2291–2304 (2014).

49. Firmino, J., Rocancourt, D., Saadaoui, M., Moreau, C. & Gros, J. Cell Division Drives Epithelial Cell Rearrangements during Gastrulation in Chick. Dev. Cell 36, 249–261 (2016).

50. Bocanegra-Moreno, L., Singh, A., Hannezo, E., Zagorski, M. & Kicheva, A. Cell cycle dynamics control fluidity of the developing mouse neuroepithelium. Nat. Phys. 19, 1050–1058 (2023).

51. Kim, S., Pochitaloff, M., Stooke-Vaughan, G. A. & Campàs, O. Embryonic tissues as active foams. Nat. Phys. 17, 859–866 (2021).

52. Kim, S. et al. A nuclear jamming transition in vertebrate organogenesis. Nat. Mater. 23, 1592–1599 (2024).

53. Ranft, J. et al. Fluidization of tissues by cell division and apoptosis. Proc. Natl. Acad. Sci. 107, 20863–20868 (2010).

54. Park, J.-A. et al. Unjamming and cell shape in the asthmatic airway epithelium. Nat. Mater. 14, 1040–1048 (2015).

55. Tetley, R. J. et al. Tissue fluidity promotes epithelial wound healing. Nat. Phys. 15, 1195–1203 (2019).

56. Saadaoui, M., Rocancourt, D., Roussel, J., Corson, F. & Gros, J. A tensile ring drives tissue flows to shape the gastrulating amniote embryo. Science 367, 453–458 (2020).

57. Niehaus, I. et al. Cerebral organoids expressing mutant actin genes reveal cellular mechanism underlying microcephaly. EMBO Rep. (2025) doi:10.1038/s44319-025-00647-7.

58. Wunderlich, S. et al. Primate iPS cells as tools for evolutionary analyses. Stem Cell Res. 12, 622–629 (2014).

59. Woltjen, K. et al. piggyBac transposition reprograms fibroblasts to induced pluripotent stem cells. Nature 458, 766–770 (2009).

60. Thevenaz, P., Ruttimann, U. E. & Unser, M. A pyramid approach to subpixel registration based on intensity. IEEE Trans. Image Process. 7, 27–41 (1998).

61. Ershov, D. et al. TrackMate 7: integrating state-of-the-art segmentation algorithms into tracking pipelines. Nat. Methods 19, 829–832 (2022).

