## Supplementary Figures S1-S5 for "Species-specific basal fluidization shapes early forebrain development"

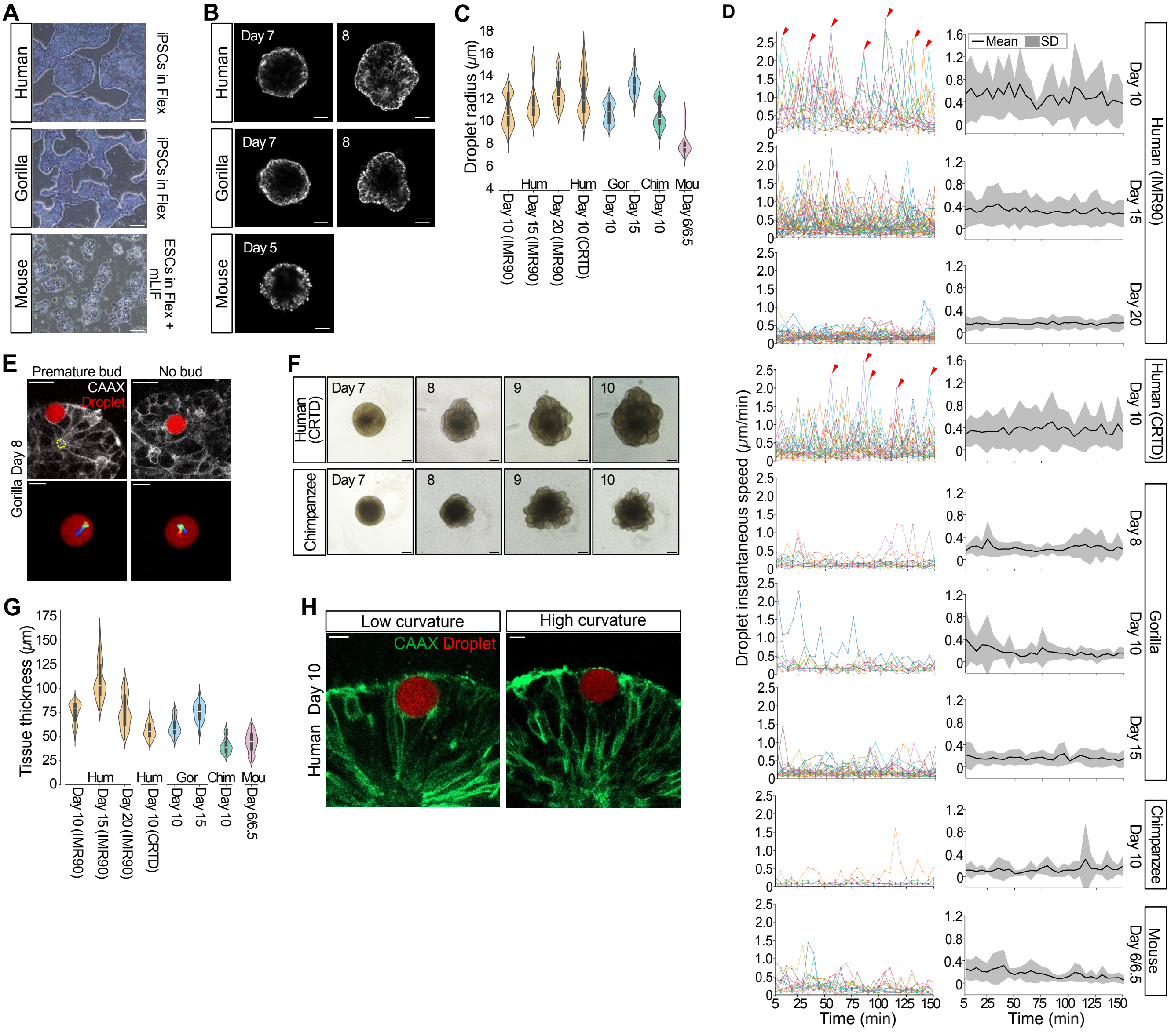
A
Human
iPSCs in Flex
Gorilla
iPSCs in Flex
Mouse
ESCs in Flex + mLIF
B
Day 7
8
Day 5
C
Droplet radius (μm)
Hum
Gor
Chim
Mou
Day 10 (IMR90)
Day 15 (IMR90)
Day 20 (IMR90)
Day 10 (CRTD)
Day 10
Day 15
Day 6/6.5
D
Mean
SD
Human (IMR90)
Human (CRTD)
Gorilla
Chimpanzee
Mouse
Day 8
Day 20
Droplet instantaneous speed (μm/min)
Time (min)
E
Premature bud
No bud
CAAX
Droplet
Gorilla Day 8
F
Human (CRTD)
Chimpanzee
9
10
G
Tissue thickness (μm)
H
Low curvature
High curvature
Human Day 10

### Fig. S1

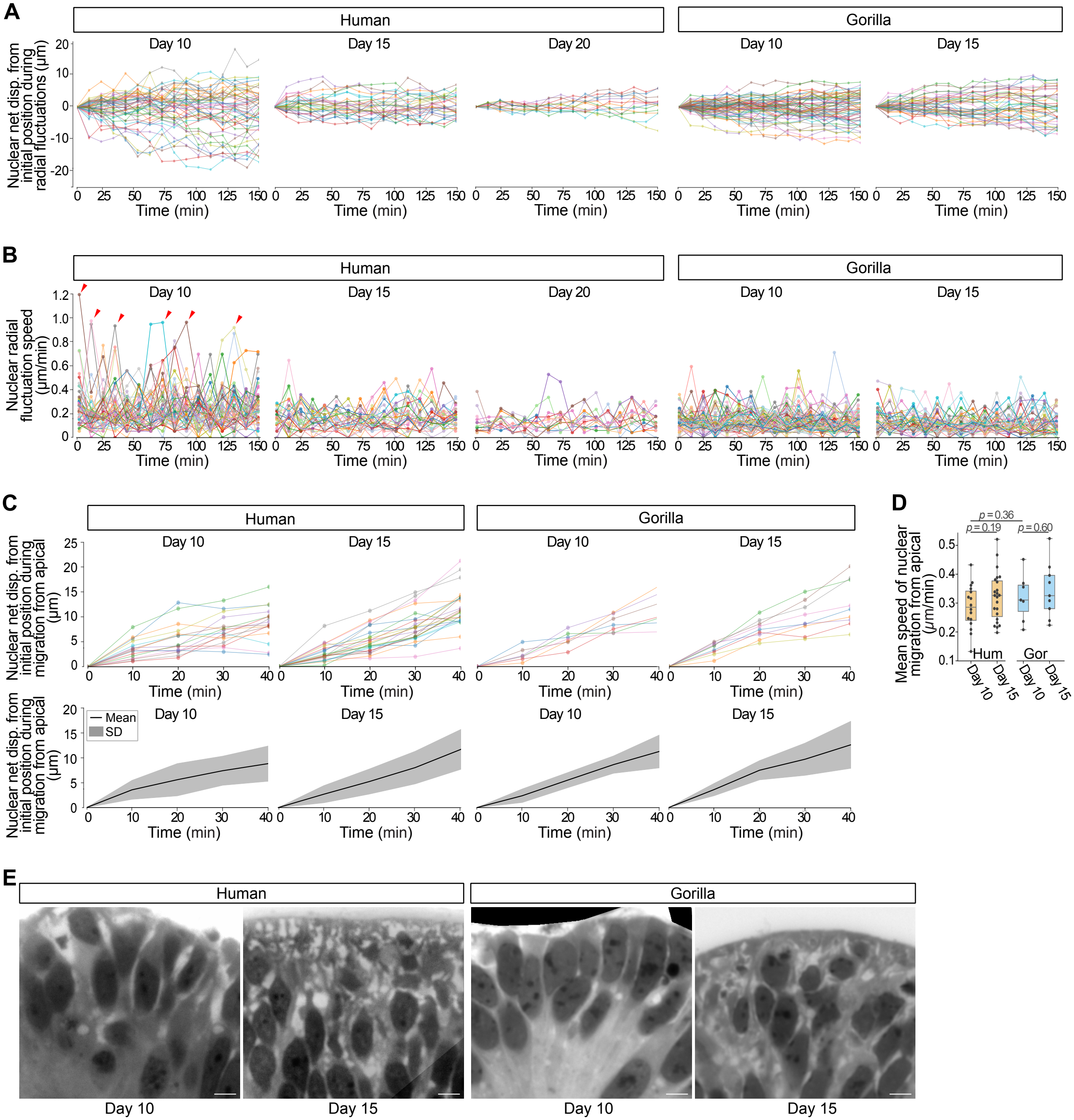
A
Human
Gorilla
Day 10
Day 15
Day 20
Nuclear net disp. from initial position during radial fluctuations (μm)
Time (min)
B
Nuclear radial fluctuation speed (μm/min)
C
Nuclear net disp. from initial position during migration from apical (μm)
Mean
SD
D
Mean speed of nuclear migration from apical (μm/min)
p = 0.36
p = 0.19
p = 0.60
Hum
Gor
E

### Fig. S2

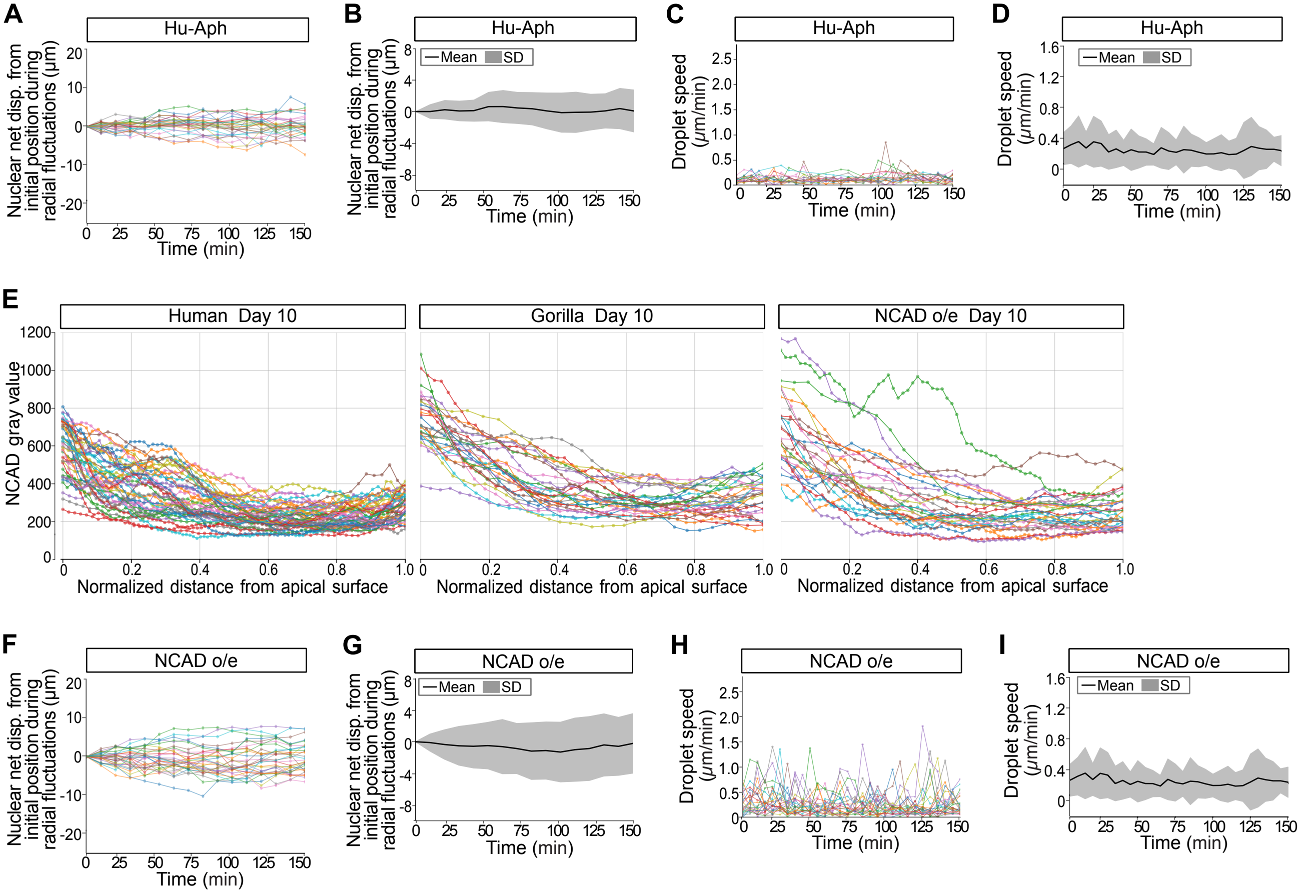
A
Hu-Aph
Nuclear net disp. from initial position during radial fluctuations (μm)
Time (min)
B
Hu-Aph
Mean
SD
Nuclear net disp. from initial position during radial fluctuations (μm)
Time (min)
C
Hu-Aph
Droplet speed (μm/min)
Time (min)
D
Hu-Aph
Mean
SD
Droplet speed (μm/min)
Time (min)
E
Human Day 10
Gorilla Day 10
NCAD o/e Day 10
NCAD gray value
Normalized distance from apical surface
F
NCAD o/e
Nuclear net disp. from initial position during radial fluctuations (μm)
Time (min)
G
NCAD o/e
Mean
SD
Nuclear net disp. from initial position during radial fluctuations (μm)
Time (min)
H
NCAD o/e
Droplet speed (μm/min)
Time (min)
I
NCAD o/e
Mean
SD
Droplet speed (μm/min)
Time (min)

**Fig. S3**

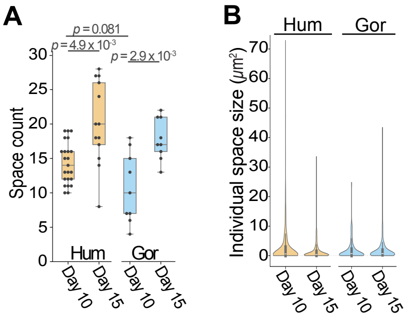


C

| Condition \ Probability | Space > 5 μm² | Space > 10 μm² | Space > 20 μm² |
|---|---|---|---|
| Human Day 10 | 0.514 | 0.244 | 0.244 |
| Human Day 15 | 0.361 | 0.093 | 0.093 |
| Gorilla Day 10 | 0.388 | 0.131 | 0.131 |
| Gorilla Day 15 | 0.374 | 0.138 | 0.138 |

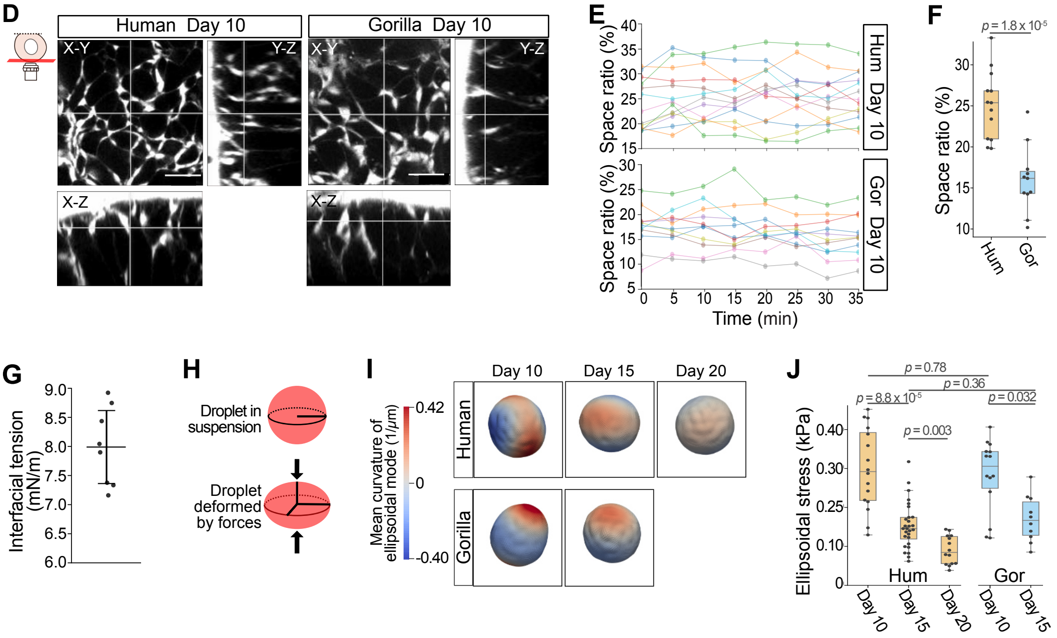


**Fig. S4**

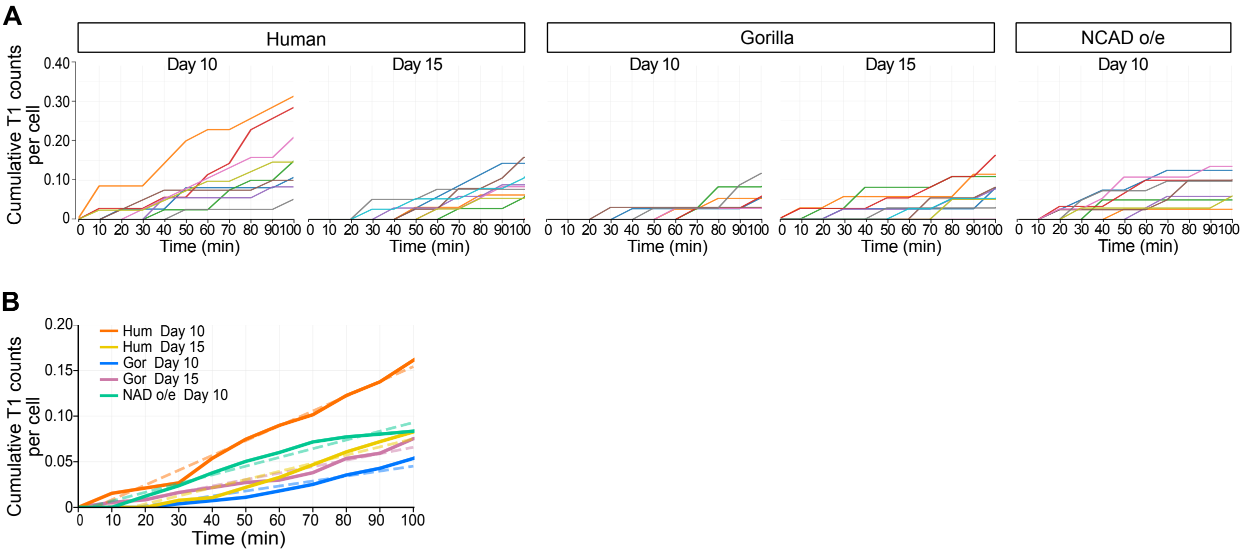

A
Human
Gorilla
NCAD o/e
Day 10
Day 15
Day 10
Day 15
Day 10
Cumulative T1 counts per cell
0.40
0.30
0.20
0.10
0
0 10 20 30 40 50 60 70 80 90 100
Time (min)
B
Hum Day 10
Hum Day 15
Gor Day 10
Gor Day 15
NAD o/e Day 10
Cumulative T1 counts per cell
0.20
0.15
0.10
0.05
0
0 10 20 30 40 50 60 70 80 90 100
Time (min)


**Fig. S5**
